# Membrane potential and feedback dynamics regulate CatSper-mediated progesterone signaling in human sperm

**DOI:** 10.1101/2025.09.14.675619

**Authors:** Michelina Kierzek, Dmitry Fridman, Cristina Biagioni, Evan W. Miller, U. Benjamin Kaupp, Christoph Brenker, Timo Strünker

**Author notes:** Contributed equally to this study (Christoph Brenker, Timo Strünker).

## Abstract

Activation of the sperm-specific Ca^2+^ channel CatSper by progesterone evokes rapid changes in intracellular Ca^2+^ in human sperm that are required for fertilization. However, the mechanisms regulating the progesterone-induced Ca^2+^ signals have remained elusive. Here, we used quantitative kinetic fluorimetry with fast voltage-sensitive fluorescent indicators to investigate how progesterone affects the membrane potential (V_m_) of human sperm. Additionally, we employed the FAST^M^ technique to simultaneously record at millisecond time resolution changes in both V_m_ and intracellular Ca^2+^. We show that progesterone evokes a rapid pulse-like depolarization and repolarization. The depolarization is caused by Ca^2+^ influx through CatSper, which pulls V_m_ away from a resting membrane potential (V_rest_) of −65 mV set by the sperm-specific K^+^ channel Slo3. We further show that V_m_- and Ca^2+^-dependent mechanisms limit the CatSper-mediated Ca^2+^ influx, thereby promoting repolarization and enabling K^+^ efflux through Slo3 channels to restore V_rest_. Our findings demonstrate that non-genomic progesterone signaling in human sperm is regulated by negative feedback on CatSper and involves a dynamic interplay between CatSper and Slo3 in controlling V_m_. We anticipate that our novel kinetic, quantitative V_m_ recording and V_m_/Ca^2+^-multiplexing techniques will reveal additional molecular mechanisms underlying CatSper-mediated Ca^2+^ signaling in human sperm both in health and disease.

## Introduction

CatSper is the principal Ca^2+^ channel in human sperm (Luo et al., 2019; Smith et al., 2013; Williams et al., 2015; Young et al., 2024), and it serves as the central hub for chemosensory signaling (Brown et al., 2019; Jeschke et al., 2021; Taiwo et al., 2021). CatSper is composed of at least fifteen subunits (reviewed by Brown et al., 2019; Hwang & Chung, 2023; Rahban & Nef, 2020; Wang et al., 2021); indeed, it is the most complex ion channel known. The channel is activated by depolarization of the membrane potential (V_m_), alkalinization of the intracellular pH (pH_i_), and by hormones such as progesterone (Jeschke et al., 2021; Lishko et al., 2011; Strünker et al., 2011). The activation of CatSper by progesterone induces a rapid, transient rise of the intracellular Ca^2+^ concentration ([Ca^2+^]_i_), followed by a slow, sustained elevation (Blackmore et al., 1990; Harper et al., 2003; Kirkman-Brown et al., 2000; Strünker et al., 2011). This Ca^2+^ signal triggers changes of the flagellar beat known as hyperactivated motility (Luo et al., 2019; Tamburrino et al., 2020; Young et al, 2024). Hyperactivation is required for human sperm to penetrate the protective egg coat and, thereby, for fertilization *in vivo* and *in vitro* (Luo et al., 2019; Young et al., 2024). Loss of CatSper function and the ensuing failure to undergo hyperactivation is a common sperm channelopathy that causes male infertility (Young et al., 2024).

CatSper is essential for sperm function, but we still do not understand how it controls the shape of Ca²⁺ signals. The transient rise of [Ca^2+^]_i_ suggests there is a complex time-dependent balance between Ca²⁺ influx and Ca²⁺ removal. Ca²⁺ is exported by plasma membrane Ca²⁺-ATPases (Herrmann et al., 2025); however, the mechanism(s) that control and limit the Ca²⁺ influx are unknown. In other cell types, like neurons, several mechanisms control Ca^2+^ flow. For example, voltage-gated Ca^2+^ channels (Ca_v_) and Ca^2+^-activated K^+^ (BK) channels work together to regulate neurotransmitter release (Berkefeld et al., 2006; Contet et al., 2016). A similar feedback mechanism may exist in human sperm, where CatSper interacts with the K^+^ channel Slo3 (*KCNU1*) (Brenker et al., 2014). Human Slo3 is a voltage-gated channel that is strongly activated by an increase of [Ca^2+^]_i_ (Brenker et al., 2014; Geng et al., 2017). When progesterone triggers Ca^2+^ influx through CatSper, this may activate Slo3, causing a hyperpolarization that could then close CatSper (Brenker et al., 2014, Kaupp & Strünker, 2017). Additionally, CatSper itself may shut off in response to changes in voltage and/or [Ca^2+^]_i_ much like other Ca_v_ channels (Budde et al., 2002). Two CatSper subunits, EFCAB9 (EF-hand calcium-binding domain-containing protein 9) and CATSPERζ, could help inactivate CatSper when [Ca^2+^]_i_ is high by stabilizing the closed channel state (Chung et al., 2017; Hwang et al., 2019).

To better understand how the progesterone-induced Ca^2+^ response is regulated in human sperm, we used fast voltage-sensitive fluorescent indicators to record the membrane potential (V_m_) of intact sperm. Moreover, we simultaneously recorded both V_m_- and Ca^2+^ signals using the multiplexing technique FAST^M^ (Kierzek et al., 2021). Our results show that non-capacitated human sperm have a resting membrane potential (V_rest_) of about −65 mV that is set by the K^+^ channel Slo3. Progesterone induces a multi-phasic V_m_ response: a rapid depolarization and repolarization, i.e., a voltage pulse, followed by a slower, smaller, and sustained increase in V_m_. We found that the initial depolarization is caused by Ca²⁺ influx through CatSper and is limited by a voltage-dependent mechanism. The following repolarization occurs because Ca^2+^ acts in a negative feedback mechanism on CatSper, reducing the rate of Ca^2+^ influx. This allows Slo3, while Ca^2+^ is flowing into the sperm, to pull V_m_ back towards V_rest_. In summary, progesterone signalling in human sperm involves complex feedback loops; these mechanisms make Ca^2+^ influx via CatSper self-limiting and ensure that the V_m_ is jointly regulated by both CatSper and Slo3.

## Results

### VoltageFluors outperform electrochromic dyes for membrane potential recordings in sperm

We studied how progesterone affects the membrane potential (V_m_) using fast voltage-sensitive fluorescent indicators. This technique was previously used to study chemosensory signaling in sea urchin sperm (Bönigk et al., 2009; Seifert et al., 2015; Trötschel et al., 2020; Strünker et al., 2006: Windler et al., 2020). There are two main types of voltage-sensitive indicators: electrochromic dyes (Loew, 2015) used for ratiometric recordings of fluorescence at two different wavelengths and photo-induced electron transfer-based indicators, or VoltageFluors (Liu & Miller, 2020), for single-wavelength recordings. To identify which indicators work best for human sperm, we first tested their performance in sea urchin sperm (*Arbacia punctulata)*, using a well-established null-point calibration method. This approach allows converting changes in fluorescence directly into millivolts (Strünker et al. 2006; Hamzeh et al., 2019). We loaded sperm with the indicators and used a stopped-flow apparatus to rapidly mix them with the chemoattractant *resact* and varying concentrations of extracellular K^+^ ([K^+^]_o_). Stimulation with resact activates the K^+^ channel CNGK (Fig. 1A), causing a hyperpolarization (Fig. 1B) that peaks at the equilibrium potential for K^+^ (E_K_) (Strünker et al., 2006). The amplitude of the hyperpolarization decreases as [K^+^]_o_ increases, ultimately reversing to a depolarization at high enough K^+^ concentrations (Fig. 1B). By plotting the maximum change in fluorescence against E_K_, we observed a linear relationship (Fig. 1C). The point where the fitted line crosses the x-axis (the ‘null-point’) (Fig. 1C, D) estimates V_rest_, while the slope of the line reflects the indicator’s performance, specifically the relative change in fluorescence (ΔF/F_0_) or fluorescence ratio (ΔR/R_0_) per millivolt (Fig. 1C, E).

**Figure 1.**
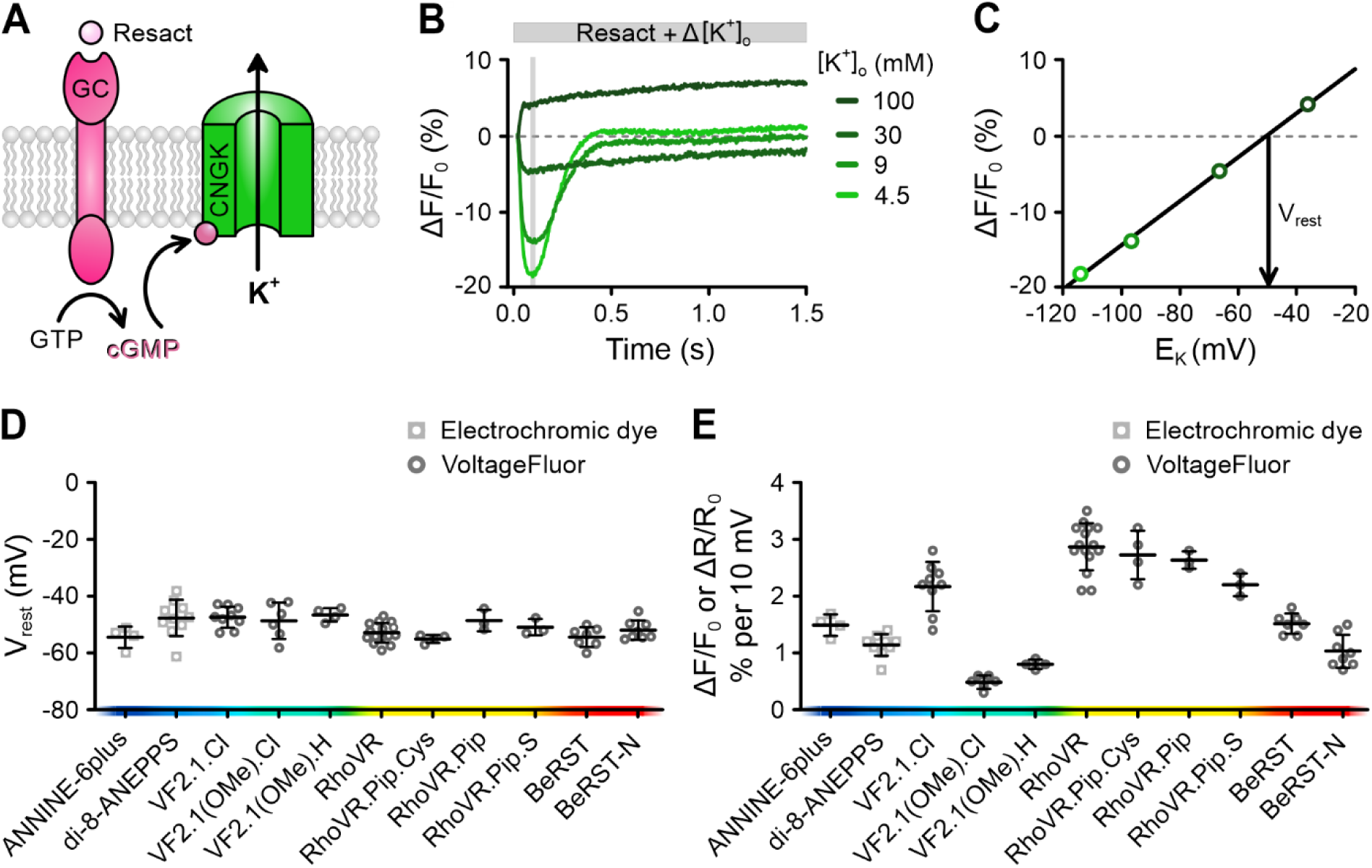
Performance of fast voltage-sensitive fluorescent indicators in sea urchin sperm. **(A)** Components of the chemosensory signaling pathway in sea urchin sperm leveraged for the null-point calibration with fluorescent voltage indicators (see explanation in text). **(B)** Representative null-point calibration using sperm loaded with the VoltageFluor RhoVR. Changes in fluorescence (ΔF/F_0_ (%)) induced by mixing sperm with buffer containing resact (2 nM) and varying concentrations of extracellular K^+^ ([K^+^]_o_), adjusting [K^+^]_o_ to the indicated values. **(C)** Relationship between the signal amplitude, averaged over the shaded window in (B), and E_K_ (calculated using [K^+^]_i_ = 423 mM). The slope of the fitted line was 2.9% (ΔF/F_0_) per 10 mV and the null-point revealed a V_rest_ of −50.5 mV. Mean (± SD) **(D)** V_rest_ and **(E)** slope determined from null-point calibrations performed using the electrochromic dyes di-8-ANEPPS and ANNINE-6plus, and various VoltageFluors (n = 3–15). Indicators are listed according to their excitation wavelength.

We compared several VoltageFluors and electrochromic dyes (Fig. 1D, E). All indicators gave similar estimates of the sea urchin sperm V_rest_ (approximately −45 mV) (Fig. 1D) (Hamzeh et al., 2019; Strünker et al., 2006; Seifert et al., 2015), but their slopes varied considerably (Fig. 1E). The best-performing electrochromic dye ANNINE-6plus (Fromherz et al., 2008) had a slope of 1.5% ΔR/R_0_ per 10 mV. However, several VoltageFluors performed even better. The top performers, VF2.1.Cl (Miller et al., 2012), the red-shifted indicator RhoVR (Deal et al., 2016) and its derivatives, RhoVR.Pip.Cys (Deal et al., 2020), RhoVR.Pip (Deal et al., 2020), and RhoVR.Pip.S (Kazemipour et al., 2019), showed slopes > 2.0% ΔF/F_0_ per 10 mV (Fig. 1E). However, some indicators such as BeRST-N (unpublished), VF2.1(OMe).Cl, and VF2.1(OMe).H (Woodford et al., 2015) were less sensitive in sea urchin sperm than in other cells (Woodford et al., 2015), demonstrating that indicator performance can vary depending on the cell type.

### Human sperm have a resting membrane potential of approximately −65 mV, which becomes less negative (depolarizes) during capacitation

To measure the V_m_ of human sperm, we used VF2.1.Cl. This indicator increases its fluorescence when the membrane depolarizes and decreases it during hyperpolarization (Miller et al., 2012). We first assessed the emission spectrum of VF2.1.Cl-loaded sperm bathed in both standard and depolarizing high-[K^+^]_o_ buffers (Fig. 2A). Depolarization led to increased fluorescence, confirming the effective integration of VF2.1.Cl into the sperm plasma membrane and reliable reporting of V_m_ changes. To determine the V_rest_ of non-capacitated sperm, we performed a null-point calibration: sperm loaded with VF2.1.Cl were mixed in a stopped-flow apparatus with buffers containing varying [K^+^]_o_ and the K^+^ ionophore valinomycin, which clamps V_m_ to E_K_. When sperm were exposed to valinomycin in standard buffer (5 mM [K^+^]_o_), fluorescence decreased, reflecting a hyperpolarization compared to V_rest_ (Fig. 2B). In contrast, upon raising [K^+^]_o_ to 24 mM or 58 mM, fluorescence increased, indicating depolarization (Fig. 2B).

**Figure 2.**
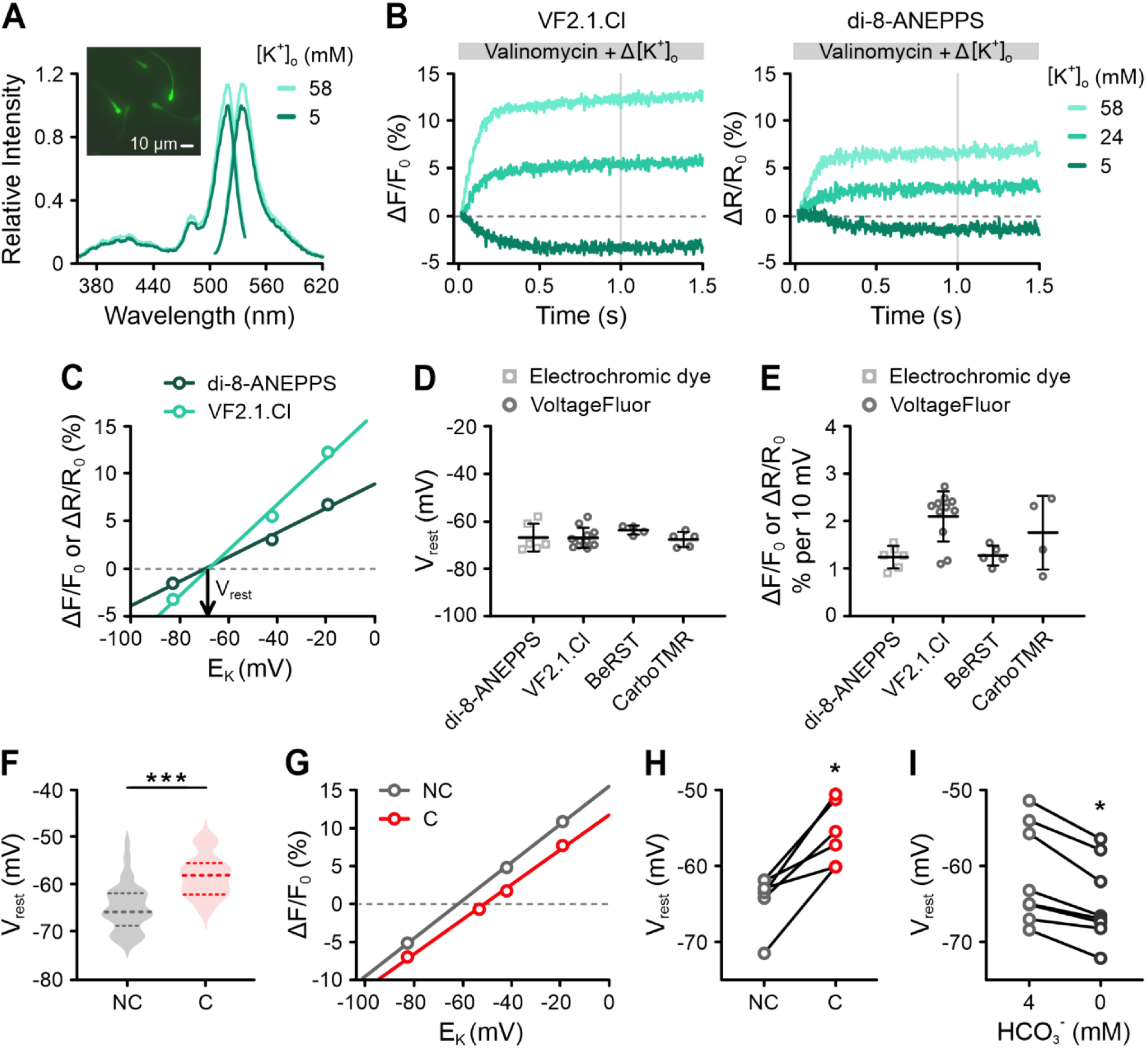
Determination of the resting membrane potential of human sperm by null-point calibration. **(A)** Excitation and emission spectra of VF2.1.Cl-loaded human sperm at 5 mM or 58 mM [K^+^]_o_. Inset: fluorescence micrograph of VF2.1.Cl-loaded sperm. **(B)** Representative changes in fluorescence (ΔF/F_0_ (%)) or the fluorescence ratio (ΔR/R_0_ (%)) from VF2.1.Cl-or di-8-ANEPPS-loaded human sperm, respectively, induced by mixing with buffer containing valinomycin and varying [K^+^]_o_, adjusting [K^+^]_o_ to the indicated values. **(C)** Relationship between the signal amplitudes, averaged over the grey windows in (B), and E_K_ (calculated using [K^+^]_i_ = 120 mM). The slopes of the fitted lines were 2.4% (ΔF/F_0_) and 1.3% (ΔR/R_0_) per 10 mV for VF2.1.Cl and di-8-ANEPPS, respectively. The x-intercepts or null-points indicated V_rest_ values of −68 mV for VF2.1.Cl-loaded sperm and −70 mV for di-8-ANEPPS-loaded sperm. **(D, E)** Mean (± SD) **(D)** V_rest_ and **(E)** slope from null-point calibrations performed using various indicators (n = 4–11). **(F)** Violin plot of the V_rest_ of non-capacitated (NC) and capacitated (C) VF2.1.Cl-loaded sperm determined by null-point calibration; the median and interquartile range are depicted as dashed lines. The V_rest_ (mean ± SD) of non-capacitated and capacitated sperm was −65 ± 5 mV (n = 110) and −58 ± 4 mV (n = 13), respectively. *** p < 0.001. **(G)** Representative paired null-point calibration using non-capacitated (NC) and capacitated (C) VF2.1.Cl-loaded sperm. The V_rest_ (x-intercept) depolarized with capacitation from −62 mV to −51 mV, while the slope remained unchanged, i.e., 2.5% and 2.3% (ΔF/F_0_) per 10 mV, respectively**. (H)** Paired plot of the V_rest_ of non-capacitated (NC) and capacitated (C) sperm (n = 6). * p < 0.05. **(I)** Paired plot of the V_rest_ of non-capacitated sperm bathed in 4 mM versus 0 mM HCO_3_^-^ (n = 7). * p < 0.05.

Plotting the amplitude of the fluorescence changes against E_K_ revealed a linear relationship (Fig. 2C). We calculated E_K_ using a [K^+^]_i_ of 120 mM (Fig. S1). In a representative calibration (Figure 2B and 2C), the slope of the fitted line was 2.4% ΔF/F_0_ per 10 mV and the x-intercept (null-point) indicated a V_rest_ of −68 mV. Across multiple experiments, the mean V_rest_ of non-capacitated sperm was −65 ± 5 mV (Fig. 2D; n = 11, Fig. 2F; n = 110; ‘n’ is the number of independent experiments using sperm from at least fifteen different donors). We validated these results using the electrochromic dye di-8-ANEPPS (Fig. 2B, C) and two VoltageFluors: BeRST (Huang et al., 2015) and isoCRhOMe (Gest et al., 2024) (Fig. 2D, E). Although the slope varied among indicators (Fig. 2E), all consistently reported a V_rest_ near −65 mV (Fig. 2D).

Next, exposing sperm to buffer containing albumin and 25 mM HCO_3_ ^-^, we examined the effect of capacitation on V_rest_. Capacitated sperm were slightly depolarized (V_rest_ of −58 ± 4 mV) (Fig. 2F; n = 13). To confirm this finding, we directly compared the V_rest_ of the same sperm sample under non-capacitating and capacitating conditions, which consistently demonstrated a depolarization following capacitation (Fig. 2G, H). Conversely, non-capacitated sperm exposed to HCO_3_ ^-^-free buffer became slightly more hyperpolarized than those in non-capacitating buffer with 4 mM HCO_3_ ^-^ (Fig. 2I).

### The membrane potential of human sperm is determined by Slo3

To identify which ionic conductances set V_rest_ in non-capacitated human sperm, we performed ion-substitution experiments. We tested whether V_rest_ depends on the extracellular concentrations of Cl^-^ ([Cl^-^]_o_), Ca^2+^ ([Ca^2+^]_o_), Na^+^ ([Na^+^]_o_), or [K^+^]_o_. Incubating sperm in low [Cl^-^]_o_, low [Ca^2+^]_o_, or high [Ca^2+^]_o_ had no effect on V_rest_ (Fig. 3A). However, in low [Na^+^]_o_, sperm hyperpolarized by −5 ± 2 mV (Fig. 3A; n = 8), suggesting that the membrane has a small Na^+^ permeability (P_Na_) that measurably contributes to V_rest_. In contrast, increasing [K^+^]_o_ from 5 mM to 25 mM caused a sizeable depolarization: the V_m_ shifted from −65 ± 5 mV to −38 ± 2 mV (Fig. 3A; n = 5). When we measured V_rest_ across a range of [K^+^]_o_, we found that V_rest_ steadily increased as [K^+^]_o_ rose (Fig. 3B). By fitting these data to the Goldman-Hodgkin-Katz equation, we calculated a P_K_/P_Na_ ratio of 37 (Fig. 3B). These results show that V_rest_ only slightly deviates from E_K_, meaning that V_rest_ is mainly determined by the K^+^ permeability.

**Figure 3.**
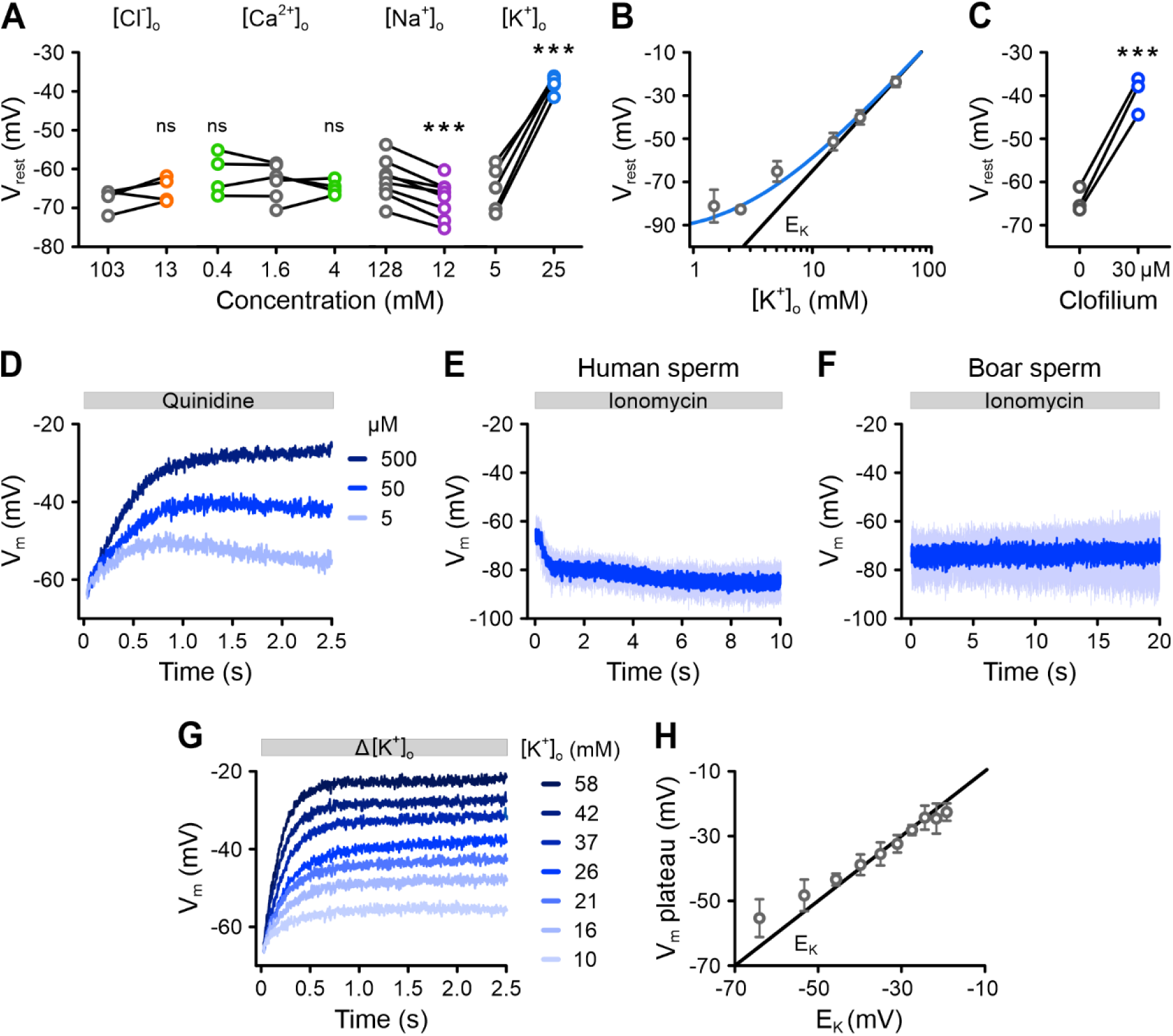
Identification of the conductances setting the resting membrane potential of human sperm. **(A)** Paired plots of the V_rest_ of non-capacitated VF2.1.Cl-loaded sperm at different concentrations of extracellular Cl^-^ ([Cl^-^]_o_), Ca^2+^ ([Ca^2+^]_o_), Na^+^ ([Na^+^]o) or K^+^, ([K^+^]o), determined by null-point calibration. *** p < 0.001. **(B)** Relationship between the mean (± SD) V_rest_ and [K^+^]_o_ (n ≥ 4). Fitting the Goldman-Hodgkin-Katz equation to the data (blue line) yielded a K^+^ versus Na^+^ permeability ratio (P_K_/P_Na_) of 37. The black line represents E_K_. **(C)** Paired plot of the V_rest_ of sperm in the absence versus presence of clofilium (30 µM) (n = 3). *** p < 0.001. **(D)** Representative changes in V_m_ induced by mixing sperm with quinidine. **(E, F)** Changes in V_m_ induced by mixing **(E)** human sperm or **(F)** boar sperm with ionomycin (10 µM) (mean ± SD; n > 3). **(G)** Average V_m_ changes induced by mixing human sperm in 5 mM [K^+^]_o_ with buffer containing increasing [K^+^]_o_, adjusting [K^+^]_o_ to the indicated values (n = 3; error bars omitted for clarity). **(H)** Relationship between E_K_ (black line) and the mean (± SD) V_m_ at the plateau of Δ[K^+^]_o_-triggered depolarizations from (G). Note that in (D), (E), (F), and (G) fluorescence was converted into millivolts by accompanying null-point calibrations (not shown).

These results agree with previous current-clamp recordings, which suggest that the V_m_ of human sperm is controlled by Slo3 channels (Brenker et al., 2014; Geng et al., 2017). To further explore Slo3’s role, we used pharmacological inhibitors. Treating sperm with clofilium, a Slo3 inhibitor (Brenker et al., 2014), caused V_rest_ to depolarize by 25 ± 4 mV (Fig. 3C; n = 3). Quinidine, another Slo3 inhibitor (Brenker et al., 2014), also triggered a rapid, dose-dependent depolarization (Fig. 3D). When we applied the Ca^2+^ ionophore ionomycin, which activates Slo3 by raising both intracellular Ca^2+^ ([Ca^2+^]_i_) and pH (pH_i_) (Fig. S2A), sperm hyperpolarized to −85 ± 5 mV (Fig. 3E; n = 5). This effect was not observed in boar sperm (Fig. 3F), which are already hyperpolarized relative to human sperm (−73 ± 4 mV), confirming that the hyperpolarization in human sperm is a genuine V_m_ response and not an artifact arising from ionomycin’s electroneutral Ca^2+^/H^+^ exchange (Erdahl et al., 1994). Finally, we examined how quickly V_m_ responds to changes in [K^+^]_o_. Increasing [K^+^]_o_ caused a rapid depolarization that followed an exponential time course, settling in < 1 s near the new E_K_ (Fig. 3G, H).

In summary, our findings demonstrate that Slo3 channels are the primary regulators of V_rest_ in human sperm, closely linking V_m_ to E_K_ (Brenker et al., 2014; Lyon et al., 2023).

### Progesterone stimulation triggers rapid V_m_ responses in human sperm

Next, we studied how progesterone affects V_m_. To investigate this, we stimulated VF2.1.Cl-loaded, non-capacitated sperm with 1 µM progesterone and recorded V_m_ (Fig. 4A). To account for variability among samples, we averaged the V_m_ responses of sperm from 38 semen samples collected from at least fifteen different donors (Fig. 4B). Progesterone consistently evoked a rapid change in V_m_. The response began with a fast depolarization, reaching −17 ± 9 mV (V_max_) in < 0.4 s (time-to-peak); this was followed by a slower repolarization phase, lasting > 1s, which brought the V_m_ to −74 ± 8 mV, which is slightly more negative than V_rest_ (Fig. 4A, B). After this initial V_m_ pulse, a slower, smaller, and sustained depolarization occurred with variable amplitude (Fig. 4A, B). A similar progesterone-induced V_m_ response was observed using the electrochromic dye di-8-ANEPPS (Fig. 4C). This V_m_ response was absent in human sperm lacking functional CatSper channels (Fig. 4D) due to a homozygous deletion of the *CATSPER2* gene (*CATSPER2*^-/-^) (Young et al., 2024). Notably, these *CATSPER2*^-/-^ sperm had a more depolarized V_rest_ (−59 ± 1 mV; n = 3) compared to control sperm (−65 ± 5 mV; *** p< 0.001). The progesterone-induced V_m_ response was also absent in boar sperm (Fig. 4E), as their CatSper channels do not respond to progesterone (Liu et al., 2021; Rehfeld et al., 2020). These results demonstrate that the progesterone-induced V_m_ response in human sperm depends on the activation of CatSper.

**Figure 4.**
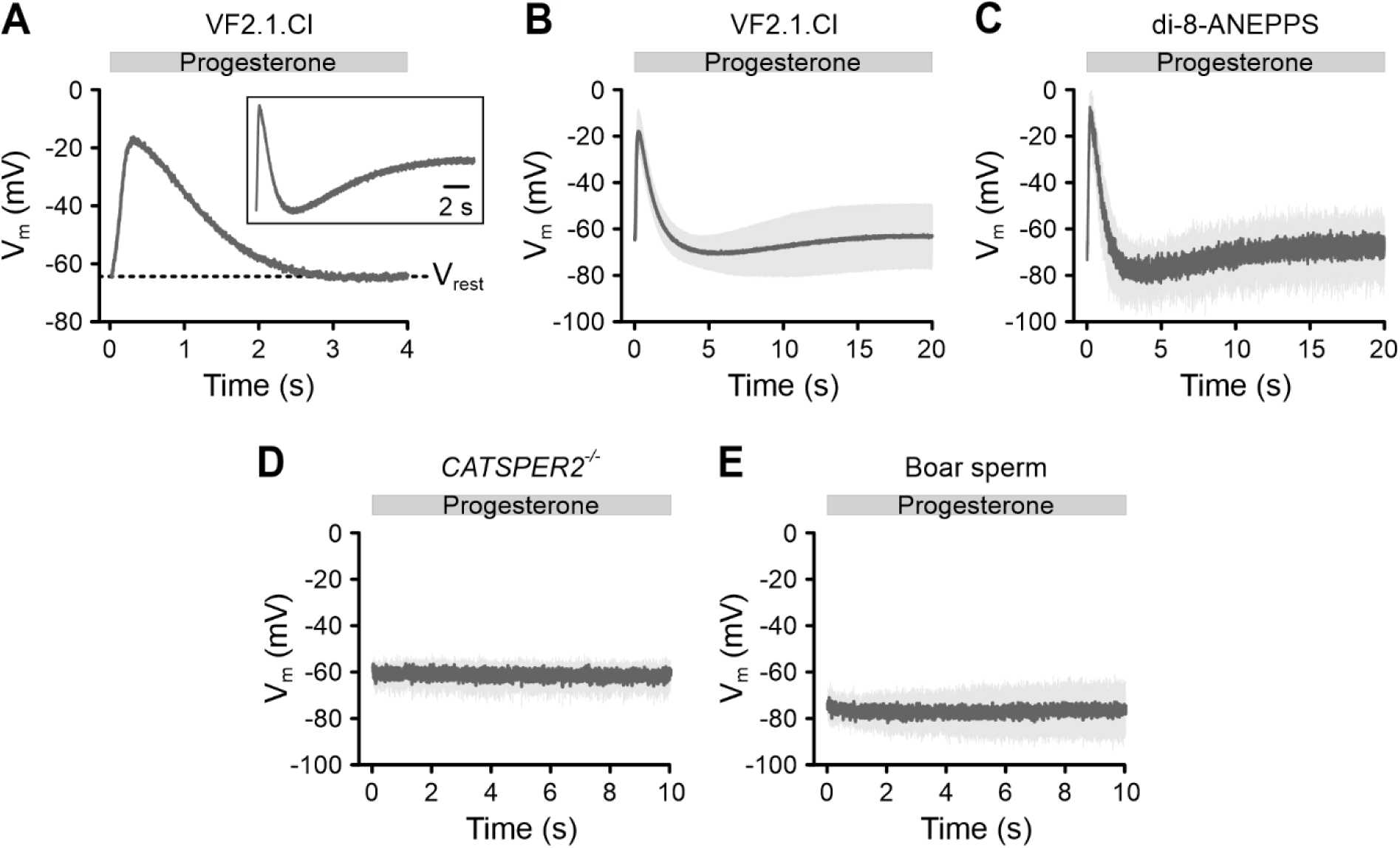
Progesterone-induced V_m_ responses of human sperm and boar sperm. **(A)** Representative progesterone-induced (1 μM) V_m_ changes in non-capacitated VF2.1.Cl-loaded human sperm. Inset: signal on an extended time scale (20 s). **(B, C)** Mean (± SD) progesterone-evoked V_m_ response in **(B)** VF2.1.Cl-loaded (n = 38) or **(C)** di-8-ANEPPS-loaded sperm (n = 3). **(D, E)** Progesterone-induced V_m_ response (mean ± SD) in VF2.1.Cl-loaded sperm from **(D)** men with a homozygous deletion of the *CATSPER2* gene (*CATSPER2*^-/-^) (n = 3) or from **(E)** boars (n = 4). Fluorescence was converted into millivolts using accompanying null-point calibrations (not shown). The V_rest_ of non-capacitated *CATSPER2*^-/-^ sperm was −59 ± 1 mV (n = 3), which is depolarized relative to non-capacitated donor sperm (*** p < 0.001).

We next studied how different progesterone concentrations affect V_m_ (Fig. 5A). At concentrations ≥ 1 nM, progesterone evoked sizeable pulse-like V_m_ responses in both non-capacitated and capacitated sperm (Fig. 5A, B). At lower nanomolar concentrations, repolarization was incomplete, and V_m_ did not return to V_rest_ level before the onset of a slower, secondary depolarization. This secondary depolarization persisted at all progesterone concentrations in non-capacitated sperm but was absent in capacitated sperm at progesterone concentrations ≥ 100 nM. As the progesterone concentration increased, the amplitude of the depolarization grew, and the depolarization and repolarization phases became faster (Fig. 5A, B). The half-maximal effective concentration (EC_50_) values ranged from 38 to 350 nM (Fig. 5C-F), depending on the signal parameter analyzed. Importantly, the depolarization phase of the V_m_ pulse was much faster than the repolarization phase (Fig. 5B, G). Analysis of the V_m_ pulse on a phase plane plot (the derivative of V_m_ over time, dV/dt, versus V_m_) demonstrated that the maximal rate of depolarization was up to 10-fold faster than that of the repolarization (Fig. 5G). These observations prompted further investigation into the molecular mechanisms underlying the V_m_ pulse.

**Figure 5.**
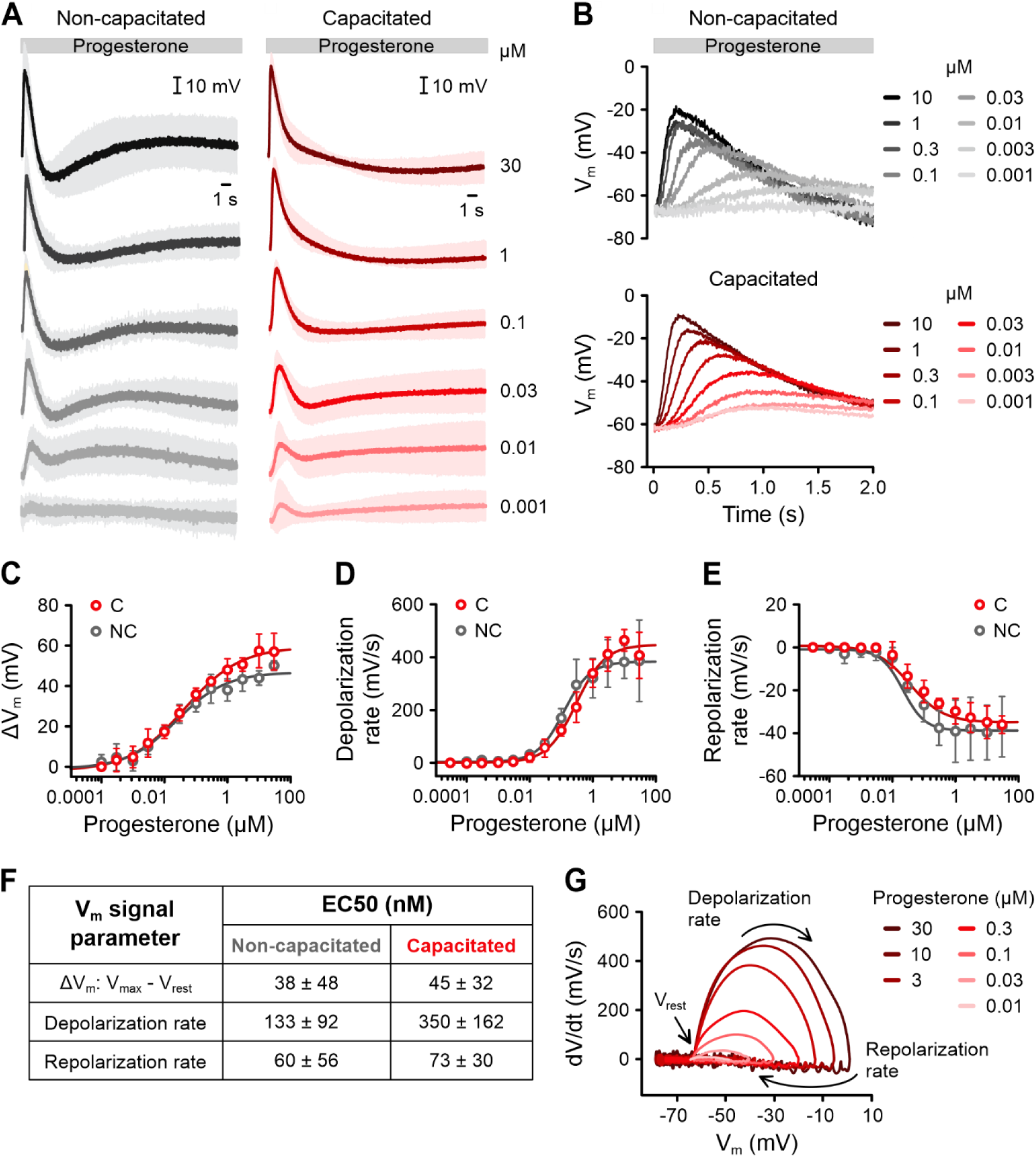
Dose-dependence of progesterone-induced V_m_ responses of human sperm. **(A)** Mean (± SD) changes in V_m_ of non-capacitated (n = 4) or capacitated (n = 3) VF2.1.Cl-loaded sperm induced by mixing with 1 nM to 30 µM progesterone. Fluorescence was converted into millivolts using accompanying null-point calibrations (not shown). **(B)** Overlay of the first 2 s of the V_m_ responses from (A). Error bars were excluded for clarity. **(C–F)** Dose-response characterization of the **(C)** amplitude (ΔV_m_ = V_max_ - V_rest_), **(D)** depolarization rate (within 40 - 60% of V_max_), and **(E)** repolarization rate (within 70 - 90% of V_max_) of the initial progesterone-induced V_m_ pulse in non-capacitated (n = 4) and capacitated sperm (n = 3). **(F)** Table with the mean (± SD) EC_50_ values. **(G)** Representative phase plane plot showing the instantaneous rate of V_m_ change (dV/dt) against V_m_ for progesterone-induced V_m_ signals from capacitated sperm.

### The repolarization phase of the V_m_ pulse does not involve Na^+^ or Cl^-^ channels

The rapid depolarization triggered by progesterone is driven by Ca^2+^ influx through CatSper. The subsequent repolarization could involve K^+^ efflux via Slo3. To clarify the role of Slo3 as well as that of Na^+^ and Cl^-^ channels in this process, we studied the progesterone-induced V_m_ pulse under varying [K^+^]_o_, at low [Na^+^]_o_, or low [Cl^-^]_o_.

Adjusting [K^+^]_o_ alters the electrochemical driving force for K^+^, which could influence the repolarization kinetics. As [K^+^]_o_ increased, the V_m_ pulse began from a more depolarized V_rest_ (Fig. 6A) and showed a smaller signal amplitude (ΔV_m_) (Fig. 6C); the rate of depolarization was also slightly affected (Fig. 6A, D). However, the rate of repolarization remained unchanged across different [K^+^]_o_ (Fig. 6A, E). Notably, higher [K^+^]_o_ caused the repolarization to dip to V_m_ values more negative than V_rest_ and even E_K_ (Fig. 6A), indicating that changes in [K^+^]_o_ have multiple effects on the V_m_ pulse. These observations make it difficult to draw definitive conclusions about the specific contribution of K^+^ efflux through Slo3 to repolarization.

**Figure 6.**
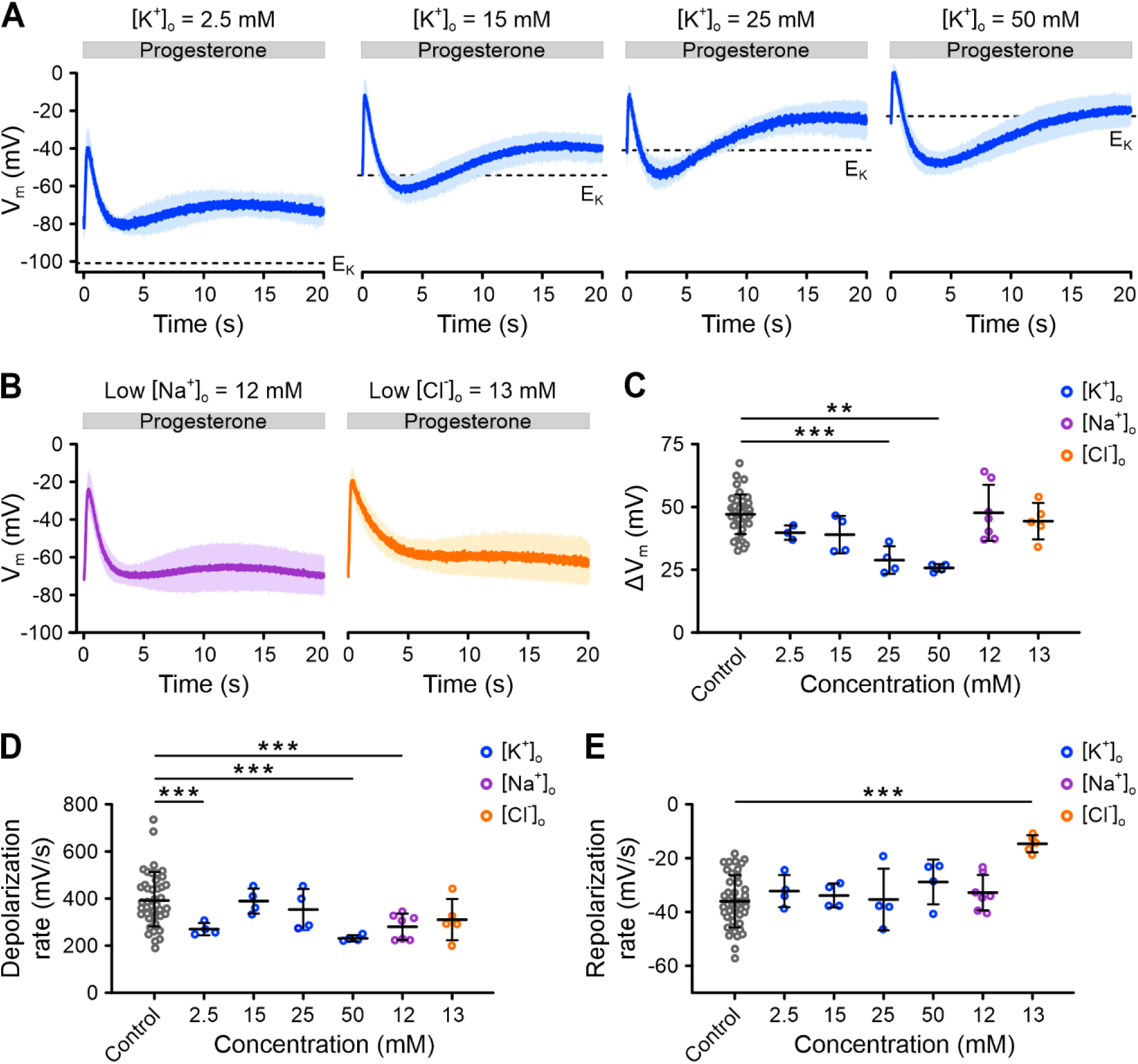
Progesterone-induced V_m_ responses of human sperm at varying extracellular K^+^, Na^+^, or Cl^-^. **(A)** Progesterone-induced (1 µM) V_m_ response (mean ± SD; n = 4) in non-capacitated VF2.1.Cl-loaded sperm at different extracellular K^+^ concentrations ([K^+^]_o_). Dashed lines indicate E_K_. **(B)** Progesterone-induced (1 µM) V_m_ response (mean ± SD) in non-capacitated, VF2.1.Cl-loaded sperm at low extracellular concentrations of Na^+^ ([Na^+^]_o_) (n = 6) or Cl^-^ ([Cl^-^]_o_) (n = 5). In (A) and (B), fluorescence was converted into millivolts using accompanying null-point calibrations (not shown). **(C–E)** Mean ± SD **(C)** amplitude (ΔV_m_ = V_max_ - V_rest_), **(D)** depolarization rate, or **(E)** repolarization rate of progesterone-induced V_m_ signals shown in (A), (B), and Figure 4B. Statistical comparisons were performed relative to the control condition (n = 38). ** p < 0.01 *** p < 0.001.

In contrast, at low (12 mM) [Na^+^]_o_, the V_m_ pulse was largely similar compared to control conditions, demonstrating that Na^+^ channels play no or only a negligible role (Fig. 6B, D, E). Finally, when we reduced [Cl^-^]_o_ to 13 mM – which shifted the E_Cl_ to approximately +30 mV, assuming an intracellular Cl^-^ concentration of 40 mM (Garcia & Meizel, 1999) – the repolarization slowed down considerably (Fig. 6B, C, E). Despite this, V_m_ still returned to the original V_rest_ of −65 mV (Fig. 6B, C, E), indicating that Cl^-^ channels are not involved in repolarization.

Given that neither Na^+^ nor Cl^-^ channels contribute to repolarization, we propose that, under physiological ionic conditions, the progesterone-induced V_m_ pulse predominantly results from Ca^2+^ influx through CatSper and K^+^ efflux through Slo3. However, additional mechanisms, such as electrogenic transport may also play a role in repolarization.

### The progesterone-induced V_m_ response is shaped by Ca^2+^- and V_m_-dependent mechanisms

We next investigated how the Ca^2+^ influx triggered by progesterone influences the repolarization phase. To modulate Ca^2+^ influx, we exposed sperm to progesterone in solutions containing different concentrations of extracellular Ca^2+^ ([Ca^2+^]_o_) and measured the V_m_ pulse. The [Ca^2+^]_o_ was adjusted by mixing sperm with a progesterone solution containing different [Ca^2+^]_o_ and/or the Ca^2+^-chelator BAPTA, resulting in a final [Ca^2+^]_o_ between 150 µM and 25 mM. We avoided using Ca^2+^ concentrations below 150 µM, as these alone elicited rapid depolarizations due to Na^+^ influx through CatSper (Fig. S3A, B) (Torres-Flores et al., 2011). Our results show that the progesterone-evoked V_m_ pulse was shaped by [Ca^2+^]_o_. The depolarization became faster as [Ca^2+^]_o_ increased, confirming that Ca^2+^ influx drives this process (Fig. 7A, C). Specifically, the rate of depolarization increased fivefold from 215 ± 50 to 1150 ± 280 mV/s between the lowest and highest [Ca^2+^]_o_ tested (Fig. 7E). Similarly, the repolarization also became faster with higher [Ca^2+^]_o_, increasing the rate of repolarization sixfold from −13 ± 4 to −77 ± 33 mV/s across the tested range (Fig. 7F). When plotted on a log-log scale, both depolarization and repolarization rates increased linearly with [Ca^2+^]_o_ (Fig. 7G), indicating a similar power-law relationship with [Ca^2+^]_o_. Doubling [Ca^2+^]_o_ sped up depolarization and repolarization by 1.23-fold. This suggests that repolarization is directly linked to Ca^2+^ influx. Interestingly, the V_m_ signal peak did not change with varying [Ca^2+^]_o_ (Fig. S4A). Instead, with increasing [Ca^2+^]_o_, the depolarization peaked more quickly (Fig. S4B), but the V_m_ at which the repolarisation began (V_max_) remained unchanged. This suggests that, while Ca^2+^ influx controls the speed of the V_m_ response, a voltage-dependent mechanism limits how far V_m_ can depolarize.

**Figure 7.**
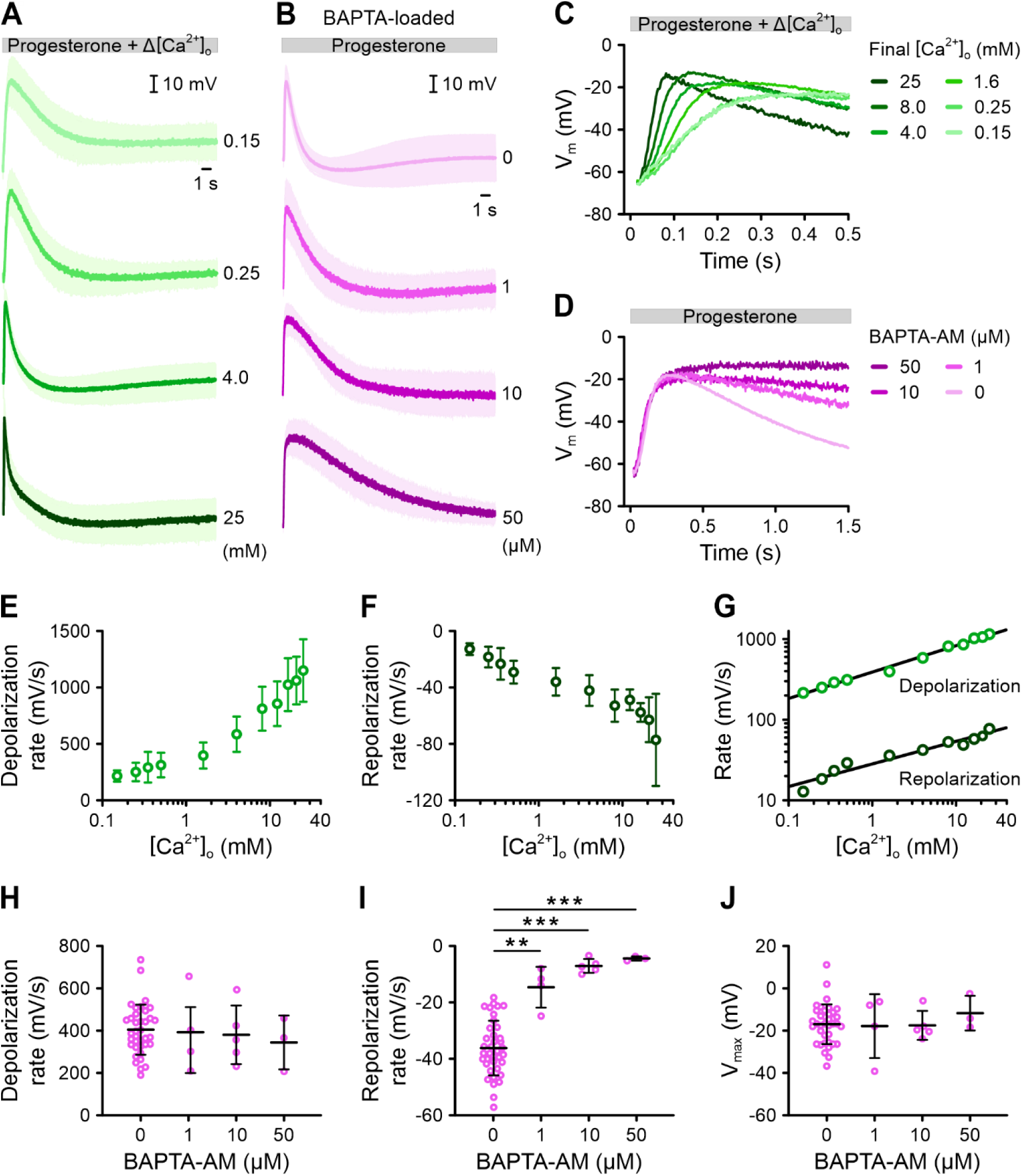
The action of extracellular Ca^2+^ and buffering of intracellular Ca^2+^ by BAPTA on progesterone-induced V_m_ responses of human sperm. **(A)** Mean (± SD) changes in V_m_ of non-capacitated VF2.1.Cl-loaded sperm induced by mixing with progesterone (1 µM) and simultaneously changing the extracellular Ca^2+^ concentration from 1.6 mM to 0.15 - 25 mM (n ≥ 4). **(B)** Mean (± SD) V_m_ changes induced by progesterone (1 µM) in sperm either pre-loaded for 60 min with 1 µM (n = 4), 10 µM (n = 5), or 50 µM (n = 3) BAPTA-AM or without, indicated as 0 μM (control; n = 38). In (A) and (B), fluorescence was converted into millivolts using accompanying null-point calibrations (not shown). **(C)** Overlay of the first 0.5 s of the V_m_ responses from (A). Error bars were excluded for clarity. **(D)** Overlay of the first 1.5 s of the V_m_ responses from (B). Error bars were excluded for clarity. **(E, F)** The mean (± SD) **(E)** depolarization rate and **(F)** repolarization rate of the V_m_ responses shown in (A) and (C) as a function of [Ca^2+^]_o_ (n ≥ 4). **(G)** Double logarithmic plot of the depolarization rate and repolarization rate against [Ca^2+^]_o_. The slopes of the fitted lines were 0.31 and 0.27, respectively. **(H, I)** The mean (± SD) **(H)** depolarization rate and **(I)** repolarization rate of progesterone-induced V_m_ responses upon BAPTA-AM loading, shown in (B) and (D) (n ≥ 3). **(J)** The mean (± SD) peak V_m_ (V_max_) reached during progesterone-induced V_m_ response upon BAPTA-AM loading, shown in (B) and (D).

To further clarify the roles of Ca^2+^ and V_m_, we loaded sperm with the Ca^2+^-chelator BAPTA to buffer intracellular Ca^2+^ (Fig. 7B, D). BAPTA neither affected V_rest_ (Fig. S3C) nor the rate of depolarization (Fig. 7H), demonstrating that intracellular BAPTA does not affect the Ca^2+^ influx. However, it significantly slowed repolarization in a dose-dependent manner (Fig. 6D, I); at the highest BAPTA loading concentration, the repolarization rate dropped almost 10-fold from −36 ± 10 mV/s to −4.5 ± 0.8 mV/s (Fig. 7B, D, I). Thus, buffering of intracellular Ca^2+^ greatly impairs repolarization, indicating that it is controlled by the inflowing Ca^2+^ and/or the resulting increase in [Ca^2+^]_i_. Despite this, the V_m_ at which repolarization started (V_max_) remained unchanged even at the highest BAPTA concentration. (Fig. 7J). This supports the idea that while Ca^2+^ influx drives depolarization, it is capped by voltage rather than Ca^2+^.

### The repolarization of V_m_ involves Ca^2+^-regulated negative feedback on CatSper

To clarify the role of Ca^2+^ in the repolarization phase, we simultaneously recorded progesterone-induced V_m_ and Ca^2+^ signals using the multiplexing technique FAST^M^ (Kierzek et al., 2021). Sperm were loaded with VF2.1.Cl and the red-shifted, low-affinity Ca^2+^ indicator Calbryte 630. We minimized the amount of Calbryte 630 loading to ensure that Ca^2+^ binding by the indicator only slightly affected the V_m_ signal (Fig. S5).

We recorded progesterone-induced V_m_ and Ca^2+^ signals at [Ca^2+^]_o_ of 1.6 mM and 8.0 mM (Fig. 8A). The depolarization and Ca^2+^ rise commenced simultaneously, without a measurable latency (Fig. 8B). However, the depolarization occurred much faster than the rise in [Ca^2+^]_i,_ and repolarization began well before the Ca^2+^ signal reached its peak (Fig. 8A). Specifically, when the repolarization started, the Ca^2+^ signal had achieved < 20% of its peak amplitude (Fig. 8C). By the time the Ca^2+^ signal reached its peak, the repolarization was already complete (Fig 8D). Increasing [Ca^2+^]_o_ from 1.6 to 8.0 mM amplified the Ca^2+^ signal peak, at which, V_m_ was more negative (Fig. 8C, D). These results show that Ca^2+^ influx occurs almost entirely during repolarization rather than during depolarization.

**Figure 8.**
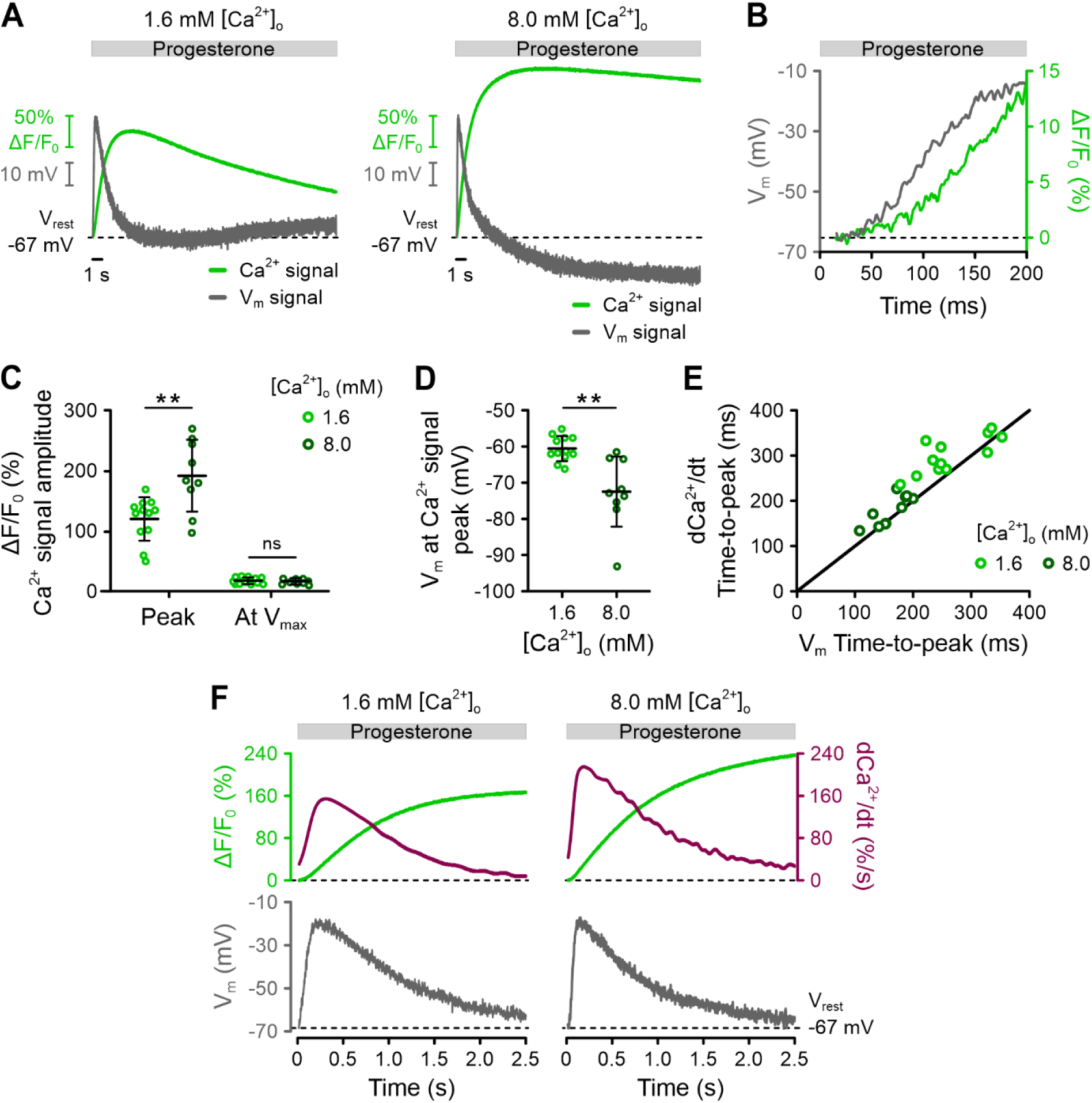
Simultaneous recording of progesterone-induced V_m_ and Ca^2+^ signals in human sperm and analysis of their temporal relationship. **(A)** Representative progesterone-evoked (1 μM) V_m_ and Ca^2+^ signals, recorded simultaneously with FAST^M^ from non-capacitated human sperm loaded with VF2.1.Cl and Calbryte 630 at 1.8 mM and 8.0 mM [Ca^2+^]_o_. **(B)** First 200 ms of V_m_ and Ca^2+^ signals from (A) at 1.6 mM [Ca^2+^]_o_. **(C)** Mean (± SD) amplitude of the progesterone-evoked (1 μM) Ca^2+^ signal at its peak and at the time when V_m_ reached V_max_, measured at 1.6 mM [Ca^2+^]_o_ (n = 12) or 8.0 mM (n = 9). ** p < 0.01. **(D)** Mean (± SD) V_m_ at the time the progesterone-evoked (1 μM) Ca^2+^ signal peaked, measured at 1.6 mM [Ca^2+^]_o_ (n = 12) or 8.0 mM [Ca^2+^]_o_ (n = 9). ** p < 0.01. **(E)** Plot showing the temporal correlation between the time-to-peak of dCa^2+^/dt and the time-to-peak of the V_m_ signal (r = 0.9074; p < 0.001). Line depicts theoretical perfect alignment. **(F)** First 3 s of the V_m_ and Ca^2+^ signals from (A) overlaid with the derivative of the Ca^2+^ signal (dCa^2+^/dt).

To gain deeper mechanistic insights into the dynamic relationship between V_m_ and [Ca^2+^]_i_, we analyzed the time derivative (dCa^2+^/dt) of the Ca^2+^ signal. Notably, the temporal pattern of dCa^2+^/dt closely mirrored that of the V_m_ pulse (Fig. 8E, F). This finding indicates that the V_m_ pulse primarily depends on the opening and closing of CatSper channels: the initial upstroke of dCa^2+^/dt reflects a rapid increase in the rate of Ca^2+^ influx as CatSper channels activate, leading to depolarization. As the rate of Ca^2+^-influx saturates, depolarization reaches its peak; after which, the rate of Ca^2+^-influx decreases due to a gradual decline in CatSper activity. During this Ca^2+^ influx, K^+^ efflux through Slo3 acts to repolarize V_m_ back toward its resting value. The decline in CatSper activity – and, thus, the kinetics of repolarization – is regulated by negative Ca^2+^ feedback, likely acting directly on CatSper.

In summary, the progesterone-induced V_m_ signal reflects the activity of CatSper channels, characterized by their rapid opening and slower, Ca^2+^-dependent closing. This dynamic creates a ‘tug-of-war’ between CatSper and Slo3 channels in regulating V_m_.

## Discussion

Reports on the V_rest_ of human sperm have varied widely, with values for non-capacitated sperm ranging from −17.7 mV (Brown et al., 2016) to −75 mV (Calzada & Telez, 1997) and from −22.7 mV (Brown et al., 2016) to −74.5 mV (Baro Graf et al., 2019) for capacitated sperm (Table S1). Even studies using the same techniques have reported inconsistent results. For instance, flow cytometry studies using the same solvatochromic voltage indicator reported the V_rest_ of non-capacitated sperm as −35.7 mV (Molina et al., 2020) and −67 mV (Matamoras-Volante et al., 2020). Additionally, the direction of capacitation-induced changes in V_rest_ have been reported as hyperpolarization (Matamoras-Volante et al., 2020; Baro Graf et al., 2020; Brown et al., 2016; Molina et al., 2020), depolarization (Calzada et al., 1988; Baro-Graf et al., 2020), or no change at all (Baro Graf et al., 2020; Matamoras-Volante et al., 2020) (Table S1). These discrepancies cannot be explained by sample variability, as many studies report average values with small standard errors (Table S1). Most of these studies used solvatochromic dyes because of their large changes in fluorescence per mV. However, as these dyes are highly sensitive to their environment, the variability in V_rest_ values (Table S1) may be due to variations in experimental conditions (dye concentration, dye-to-sperm ratio, pH, absence/presence of albumin, etc.) and/or technical difficulties (Hladky and Rink, 1976; Tsien and Hladky, 1978; Plasek and Hrouda, 1991; Zeng et al., 1995). Taken together, this suggests that while solvatochromic dyes are suitable for qualitative reporting of slow, voltage-dependent fluorescence changes under defined conditions (Luque et al., 2023), they are less suitable for quantitative determinations of V_rest_ in human sperm. Therefore, we employed fast electrochromic dyes and VoltageFluors, which, combined with quantitative kinetic fluorimetry using the stopped-flow technique, have been instrumental and reliable in determining V_rest_ and in delineating the chemosensory signaling pathway in sea urchin sperm (Bönigk et al., 2009; Hamzeh et al., 2019; Seifert et al., 2015; Strünker et al., 2006; Trötschel et al., 2020; Windler et al., 2020). Here, we adapted this approach to human sperm to investigate how V_rest_ is regulated. Our findings show that in human sperm, V_rest_ is controlled by Slo3 K^+^ channels, consistent with earlier research (Brenker et al., 2014; Lyon et al., 2023). We observed that non-capacitated human sperm have a V_rest_ of −65 mV, which becomes slightly less negative during capacitation. This observation contrasts with previous dogma suggesting that non-capacitated sperm are rather depolarized and that V_rest_ hyperpolarizes significantly upon capacitation, thereby regulating key physiological processes involved in sperm maturation and fertilization (Ritagliati et al., 2018). Based on our comprehensive study of V_rest_, we propose that a capacitation-induced hyperpolarization is, in fact, lacking in human sperm.

In contrast, capacitation-induced hyperpolarization is well-established in mouse sperm. Across various methods, non-capacitated mouse sperm consistently show a V_rest_ of approximately −40 mV, which hyperpolarizes to around −60 mV during capacitation (Table S2). This hyperpolarization is mediated by Slo3 (Santi et al., 2010; Zeng et al., 2011) and is essential for preparing mouse sperm for hyperactivation and acrosomal exocytosis (De La Vega-Beltran et al., 2012; Zeng et al., 1995; Santi et al., 2010; Zeng et al., 2011). However, caution is needed when extrapolating these findings from mice to humans as the two species differ in their ion channels and transporters in the membrane. For instance, mouse Slo3 channels are gated by pH_i_ (Santi et al., 2010; Zeng et al., 2011), whereas in humans, Slo3 is primarily regulated by [Ca^2+^]_i_ (Brenker et al., 2014; Geng et al., 2017). Based on our analysis, we propose that V_rest_ and its control by Slo3 differ between mouse and human sperm, and that capacitation-induced hyperpolarization is not a feature of human sperm. This distinction highlights the importance of species-specific studies when investigating the physiology of sperm function (Kaupp & Strünker, 2017).

We further show that progesterone triggers a rapid V_m_ pulse, reflecting dynamic Ca^2+^ influx through CatSper. This influx is regulated by both voltage- and Ca^2+^-dependent feedback mechanisms. The initial depolarization results from Ca^2+^ influx via CatSper, shifting V_m_ away from the Slo3-determined V_rest_. A voltage-dependent mechanism limits this depolarization.

Repolarization occurs when CatSper activity declines, which likely involves Ca^2+^-dependent inactivation, reducing the rate of Ca^2+^ influx and allowing K^+^ efflux through Slo3 channels to restore V_rest_. The inactivation kinetics probably determine the amplitude and shape of the Ca^2+^ signal.

The negative voltage- and Ca^2+^-dependent feedback on CatSper may involve Ca^2+^-binding proteins such as the CatSper subunits EFCAB9-CATSPERζ or other CatSper-associated proteins like CaMKII or calcineurin (Chung et al., 2014). To identify the specific molecular mediator, future studies should examine progesterone-induced V_m_ responses in the presence of selective inhibitors, such as cyclosporin (Miyata et al., 2015). Although pharmacological tools targeting EFCAB9-CATSPERζ are currently unavailable, biochemical experiments using recombinant human EFCAB9 and CATSPERζ could clarify the Ca^2+^-and pH-dependence of their interaction, as demonstrated in mouse studies (Hwang et al., 2019). If pH also regulates the V_m_ response – potentially by modulating EFCAB9-CATSPERζ-mediated Ca^2+^-dependent inactivation – simultaneous monitoring of V_m_, [Ca^2+^]_i_, and pH_i_ using FAST^M^ technology could reveal how these factors interact to shape the sperm’s response to progesterone.

While we could not definitively establish Slo3’s role in the progesterone-induced V_m_ response, it might involve activation of Slo3 by Ca^2+^. Slo3 may become more activated after the initial V_m_ pulse, coinciding with the peak of the Ca^2+^ signal. Supporting this idea, V_m_ was more hyperpolarized when the Ca^2+^ signal peak was higher (Fig. 8E). Therefore, additional studies are needed to clarify Slo3’s function in the response to progesterone, such as examining the V_m_ response in the presence of Slo3 inhibitors (Lyon et al., 2023) or using sperm from infertile men lacking functional Slo3 (Lv et al., 2022). Downstream activation of Slo3 may complement the negative feedback on CatSper, acting as a secondary ‘break’ that limits CatSper activation or facilitates its recovery from inactivation, shaping CatSper-mediated Ca^2+^ signals.

CatSper’s activation by progesterone is self-limiting, due to its control by negative feedback mechanisms. This mechanism might allow CatSper to perform specific functions while protecting sperm from harmful Ca^2+^ overload. Excessive [Ca^2+^]_i_ can halt sperm movement (Okunade et al., 2004; Schuh et al., 2004; Tateno et al., 2013), promote the degradation of CatSper proteins (Chung et al., 2014; Ded et al., 2020), and trigger the exposure of phosphatidylserine at the outer membrane leaflet (Martin et al., 2005), a signal that marks sperm for removal by phagocytic immune cells in the female reproductive tract. If the negative feedback fails, it might cause forms of CatSper-related male infertility that current tests cannot detect (Young et al., 2024). Targeting the feedback control of CatSper could also be an approach for contraception. While current strategies aim to block CatSper, small molecules that cause uncontrolled CatSper activation – and thus toxic Ca^2+^ overload – might also be effective at preventing fertilization.

Our optical recording technique has revealed Ca^2+^-dependent mechanisms controlling CatSper that cannot be directly studied with traditional electrophysiology. Ca^2+^ currents carried by CatSper are too small to be measured directly, so monovalent currents are recorded instead (Liu et al., 2021, Lishko et al., 2011, Strünker et al., 2011). However, these conditions are not physiological as they lack Ca^2+^. For example, progesterone produces a transient Ca^2+^ signal recorded with fluorimetry, whereas monovalent progesterone-evoked CatSper currents recorded with electrophysiology do not inactivate (Lishko et al., 2011, Strünker et al., 2011). The minimally invasive optical technique showcased here has uncovered new aspects of CatSper function and holds promise for further discoveries about signaling in human sperm. CatSper acts as a polymodal sensor, responding to a variety of internal and external factors (Brenker et al., 2012; Jeschke et al., 2021; Schiffer et al., 2014; Strünker et al., 2011). Measuring changes in V_m_ provides a new way to distinguish between different signaling modalities, which may not be apparent from Ca^2+^ signals alone. For instance, although pregnenolone sulfate, prostaglandin E1 (PGE1), and NH_4_Cl-induced alkalinization all cause a transient rise in [Ca^2+^]_i_ (Jeschke et al., 2021; Strünker et al., 2011), their effects on V_m_ differ (Fig. 9). Pregnenolone sulfate produces a V_m_ response similar to progesterone (Fig. 8A), while V_m_ responses triggered by PGE1 and NH_4_Cl feature less pronounced repolarization phases (Fig. 9B, C). This suggests that different stimuli involve distinct feedback mechanisms and/or signaling pathways downstream of CatSper activation.

**Figure 9.**
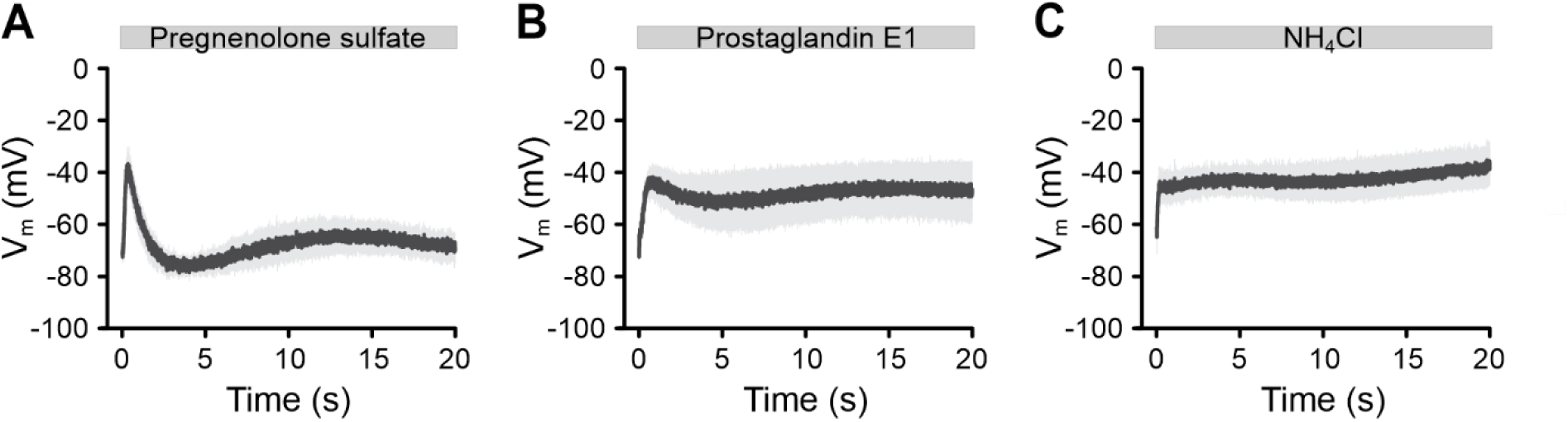
Pregnenolone sulfate-, prostaglandin E1-, or alkaline-evoked V_m_ response in human sperm. **(A–C)** Mean (± SD) changes in V_m_ of non-capacitated VF2.1.Cl-loaded sperm induced by mixing with **(A)** 1 µM pregnenolone sulfate (n = 4), **(B)** 1 µM prostaglandin E1 (n = 4), or **(C)** 30 mM NH_4_Cl (n = 4).

We anticipate that kinetic V_m_ recordings, especially when combined with FAST^M^ for multi-modal analysis, will provide deeper insights into the signaling networks involving CatSper, Slo3, and electrogenic transporters (Wang et al., 2021) that control the function of human sperm.

## Materials and Methods

### Reagents

Unless otherwise stated, reagents were obtained from Sigma-Aldrich. VoltageFluors were synthesized by the laboratory of Evan W. Miller at the University of California, Berkeley.

### Sperm preparation

Sperm collection from *Arbacia punctulata* was performed as previously described (Hamzeh et al., 2019). In brief, males were induced to spawn by injecting 500 µl of 0.5 M KCl into their body cavity; the spawn (dry sperm) was collected with a Pasteur pipette and stored on ice. Artificial sea water (ASW) and high K^+^ ASW were prepared as previously described (Hamzeh et al., 2019). Human semen samples were provided by normozoospermic donors and men with a homozygous deletion of the *CATSPER2* gene, resulting in loss of CatSper function (for details, see Schiffer et al., 2020; Young et al., 2024). All participants gave prior written consent, in accordance with protocols approved by the Ethics Committee of the Ärztekammer Westfalen-Lippe and the Medical Faculty Münster (4INie, 2021-402-f-S) as well as the Declaration of Helsinki. Semen samples were produced by masturbation into plastic containers. Following liquefaction for ≤ 30 min, motile sperm were purified from the ejaculate using the swim-up technique (Strünker et al., 2011), modified into two steps for higher yield. Notably, this swim-up method enriches motile sperm while depleting (dilution factor ≥ 15,000) diffusible molecules from the seminal plasma (e.g., Zn^2+^, prostaglandins, etc.). First, ≤ 1 ml of ejaculate was layered in a 50-ml falcon tube below 4 ml of Human Tubular Fluid (HTF) medium containing (in mM): 93.8 NaCl, 4.7 KCl, 0.2 MgSO_4_, 0.369 KH_2_PO_4_, 2.04 CaCl_2_, 0.33 Na-pyruvate, 20.98 HEPES, 2.78 glucose, 21.4 Na-lactate, 4 NaHCO_3_, pH 7.35 at 37°C (adjusted with NaOH). After incubation for 1 hr at 37°C, the uppermost 3 ml of HTF containing motile sperm was aspirated and transferred to 15-ml conical tubes. To further remove seminal plasma-derived molecules (washing), the sperm suspension was diluted with HTF to 15 ml, centrifuged (20 min, 700 g, 37°C), and the supernatant was discarded. The sperm pellet was resuspended in 15 ml of HTF, centrifuged again (20 min, 700 g, 37°C), and the supernatant was discarded. Finally, the sperm pellet was resuspended in HTF to a density of 1 x 10^7^ sperm/ml and kept at 37°C. Next, the leftovers from the initial swim-up, i.e., the ejaculate layer with 1 ml HTF, were collected, diluted tenfold with HTF, and centrifuged (20 min, 700 g, 37°C). Pellets were resuspended in 20 ml HTF, aliquoted in 5 ml portions into 50-ml falcon tubes, briefly centrifuged (5 min, 700 g, 37°C), and incubated for 1 hr at 37°C to further allow motile sperm to swim-up. The subsequent collection of motile sperm and washing was repeated as previously described, yielding another suspension at 1 x 10^7^ sperm/ml. The two sperm suspensions were combined and collectively referred to as swim-up sperm. Additionally, swim-up sperm from different, same-day donors were also routinely combined. For *in vitro* capacitation, the swim-up procedure was modified such that sperm pellets after the second centrifugation were resuspended in capacitating HTF medium (HTF^C^) containing (in mM): 72.8 NaCl, 4.7 KCl, 0.2 MgSO_4_, 0.369 KH_2_PO_4_, 2.04 CaCl_2_, 0.33 Na-pyruvate, 20.98 HEPES, 2.78 glucose, 21.4 Na-lactate, 25 NaHCO_3_, and 3 mg/ml human serum albumin (Irvine Scientific), pH 7.35 at 37°C (adjusted with NaOH), and incubated for at least 2 hours at 37°C in a 10% CO_2_ atmosphere. Boar sperm (*Sus scrofa domesticus*) were obtained from GFS-Genossenschaft zur Fӧrderung der Schweinehaltung eG (Ascheberg, Germany) in TALP medium, containing (in mM): 90 NaCl, 3 KCl, 0.4 MgCl_2_, 0.3 Na_2_HPO_4_, 2 CaCl_2_, 2 Na-pyruvate, 10 HEPES, 5 glucose, 21.6 Na-lactate, 25 NaHCO_3_ and 3 mg/ml bovine serum albumin. Sperm were diluted fivefold in HTF, centrifuged (20 min, 700 g, 37°C), and resuspended in HTF to a density of 1 x 10^7^ sperm/ml and kept at 37°C.

### Determination of the intracellular K^+^ concentration in human sperm

At least 3.4 x 10^7^ swim-up sperm in HTF were centrifuged (20 min, 700 g, 37°C) after being diluted with K^+^-free HTF medium containing (in mM): 98.5 NaCl, 0.2 MgSO_4_, 2.04 CaCl_2_, 0.33 Na-pyruvate, 20.98 HEPES, 2.78 glucose, 21.4 Na-lactate, 4 NaHCO_3_, pH 7.35 at 37°C (adjusted with NaOH). The sperm pellet was then resuspended in 100 µl K^+^-free HTF. Subsequently, 500 µl of double-distilled H_2_O supplemented with 0.001% digitonin was added to permeabilize the sperm and, thereby, release their cytosolic K^+^. The density of this permeabilized-sperm suspension was precisely determined using a Neubauer chamber (Neubauer-Improved, Assistent) and by flow cytometry (CytoFLEX S, Beckman Coulter). The sperm were then pelleted by centrifugation (10 min, 3000 g, 4°C), and 550 µl of the sperm-free supernatant was aspirated, heated for 10 min at 90°C, and its K^+^ concentration was determined by inductively coupled mass spectrometry (Xell AG). To correct for residual K^+^ remaining in the medium prior to sperm permeabilization, this procedure was performed in parallel with an equivalent volume of sperm-free medium as a background control (see schematic in Fig. S1). Taking into account the sperm density, the background control-corrected K^+^ concentration of the supernatant, and a cytoplasmic volume of 22 fl (Curry et al., 1996), we determined that sperm have an intracellular K^+^ concentration ([K^+^]_i_) of 120 ± 40 mM (Fig. S1B, C; n = 3), which is consistent with previous reports (Linares-Hernandez et al., 1998).

### Loading of sperm with fluorescent indicators

Sea urchin sperm were loaded with fluorescent voltage indicators as described before (Kierzek et al., 2021). In brief, dry sperm were diluted 1:6 (v/v) in ASW supplemented with 0.05% Pluronic F-127 and either 10 µM of a VoltageFluor or 20 µM of the electrochromic dyes di-8-ANEPPS (ThermoFisher) or ANNINE-6plus (Dr. Hinner & Dr. Hübener GbR). This sperm suspension was incubated for 5 min at 18°C in the dark, diluted 1:20 (v/v) with ASW, and allowed to equilibrate for 5 min before measurement. Boar sperm and human swim-up sperm (1 x 10^7^ sperm/ml in HTF) were loaded with either 1 µM VF2.1.Cl (Miller et al., 2012), 500 nM BeRST (Huang et al., 2015), or 500 nM isoCRhOMe (Gest et al., 2024) in the presence of 0.05% Pluronic F-127 for 10 min at 37°C, or with 10 µM di-8-ANEPPS in the presence of 0.025% Pluronic F-127 for 5 min at 37°C. After loading, the sperm suspension was diluted fivefold with HTF (pH adjusted at 30°C) and then centrifuged (10 min, 700 g, 30°C) to remove excess indicator. The indicator-loaded sperm pellets were resuspended to a density of 6 x 10^7^ sperm/ml in HTF (pH adjusted at 30°C). For human sperm in capacitating conditions (HTF^c^ for ≥ 2 hours), sperm were first diluted fivefold in HSA-free HTF^C^, centrifuged (20 min, 700 g, 37°C), resuspended in HSA-free HTF^C^ to a density of 1 x 10^7^ sperm/ml, and then loaded with VF2.1.Cl as described above. For experiments involving intracellular BAPTA, human swim-up sperm (1 x 10^7^ sperm/ml) in HTF were incubated with either 1 µM, 10 µM, or 50 µM BAPTA AM (ThermoFisher) and 0.05% Pluronic F-127 for 1 hr at 37°C. After BAPTA-loading, the sperm suspension was diluted fivefold with HTF and centrifuged (10 min, 700 g, 37°C). The sperm pellet was then resuspended in HTF to a density of 1 x 10^7^ sperm/ml and incubated for 30 min at 37°C; after which, the VF2.1.Cl-loading protocol proceeded as described above.

For FAST^M^ recordings of V_m_ and Ca^2+^ signals, sperm were loaded with 4 μM Calbryte 630 AM (AAT Bioquest) in the presence of 0.05% Pluronic F-127 for 1 min at 37°C. Excess indicator was removed by performing a sixfold dilution in HTF, followed by centrifugation (10 min, 700 g, 37°C). The Calbryte 630-loaded sperm pellets were resuspended to a density of 1 x 10^7^ sperm/ml, incubated for 30 min at 37°C to allow for complete de-esterification of Calbryte 630 AM, and then loaded with VF2.1.Cl as described above.

For recordings of pH_i_ signals, sperm were loaded with 2.5 µM of the fluorescent pH-sensitive indicator pHrodo AM (ThermoFisher) in the presence of 0.05% Pluronic F-127 for 10 min at 37°C. After loading, the sperm suspension was diluted fivefold with HTF (pH adjusted at 30°C) and centrifuged (10 min, 700 g, 30°C). The sperm pellet was resuspended to a density of 6 x 10^7^ sperm/ml in HTF (pH adjusted at 30°C).

### Experiments with human sperm at varying extracellular K^+^, Na^+^, Cl^-^, or Ca^2+^

To study the effects of [K^+^]_o_ on V_rest_ and the progesterone-induced V_m_ response, [K^+^]_o_ was adjusted during the final step of the indicator-loading protocol. After washing with standard HTF ([K^+^]_o_ = 5.06 mM), the indicator-loaded sperm pellet was resuspended in HTF containing either 1.49, 2.50, 5.06, 15.0, 25.5, or 50.2 mM [K^+^]_o_. These [K^+^]_o_-variants of HTF were prepared by mixing standard HTF in varying proportions with either K^+^-free HTF, which contained (in mM): 98.5 NaCl, 0.2 MgSO_4_, 2.04 CaCl_2_, 0.33 Na-pyruvate, 20.98 HEPES, 2.78 glucose, 21.4 Na-lactate, 4 NaHCO_3_, pH 7.35 at 30°C (adjusted with NaOH), or High-K^+^ HTF ([K^+^]_o_ = 110.37 mM), which contained (in mM): 98.5 KCl, 0.2 MgSO_4_, 0.369 KH_2_PO4, 2.04 CaCl_2_, 0.33 Na-pyruvate, 20.98 HEPES, 2.78 glucose, 21.4 Na-lactate, 4 NaHCO_3_, pH 7.35 at 30°C (adjusted with KOH). To study in a paired fashion the effects of the Slo3 inhibitor clofilium on V_rest_, VF2.1.Cl-loaded sperm at a density of 6 x 10^7^ sperm/ml in HTF (pH adjusted at 30°C) were incubated in the presence (30 µM) or absence of clofilium for 5 min at 30°C prior to measurement.

To study the V_m_ response evoked by increases in [K^+^]_o_ (Fig. 3F-H), sperm bathed in standard HTF ([K^+^]_o_ = 5.06 mM) were mixed in the stopped-flow apparatus with HTF containing 15, 26, 37, 47, 58, 68, 79, 89, 100, or 110 mM [K^+^]_o_. These [K^+^]_o_-variants of HTF were prepared by mixing standard HTF ([K^+^]_o_ = 5.06 mM) in varying proportions with High-K^+^ HTF ([K^+^]_o_ = 110.37 mM). The final [K^+^]_o_ in the cuvette after mixing was 10, 16, 21, 26, 31, 37, 42, 47, 52, or 58 mM, respectively.

To study the effects of [Na^+^]_o_ on V_rest_ and the progesterone-induced V_m_ response, [Na^+^]_o_ was adjusted during the indicator-loading protocol. After loading in standard HTF ([Na^+^]_o_ = 129.5 mM), the sperm suspension was diluted with Low-Na^+^ HTF ([Na^+^]_o_ = 4.33 mM), which contained (in mM): 115.2 N-methyl-D-glucamine (NMDG), 4.7 KCl, 0.2 MgSO_4_, 0.369 KH_2_PO4, 2.04 CaCl_2_, 0.33 Na-pyruvate, 20.98 HEPES, 2.78 glucose, 21.4 lactic acid, 4 NaHCO_3_, pH 7.35 at 30°C (adjusted with HCl). This dilution reduced [Na^+^]_o_ to 24 mM. After centrifugation (10 min, 700 g, 30°C), the sperm pellet was resuspended in Low-Na^+^ HTF, further reducing [Na^+^]_o_ to 12 mM.

To study the effects of [Cl^-^]_o_ on V_rest_ and the progesterone-induced V_m_ response, [Cl^-^]_o_ was adjusted during the indicator-loading protocol. After loading in standard HTF ([Cl^-^]_o_ = 96 mM), the sperm suspension was diluted with Low-Cl^-^ HTF ([Cl^-^]_o_ = 9.08 mM), which contained (in mM): 98.5 aspartic acid, 93.8 NaOH, 4.7 KOH, 0.2 MgSO_4_, 0.369 KH_2_PO4, 4.55 CaCl_2_, 0.33 Na-pyruvate, 20.98 HEPES, 2.78 glucose, 21.4 lactic acid, 4 NaHCO_3_, pH 7.35 at 30°C (adjusted with NaOH). This dilution reduced [Cl^-^]_o_ to 26 mM. After centrifugation (10 min, 700 g, 30°C), the sperm pellet was resuspended in Low-Cl^-^ HTF, further reducing [Cl^-^]_o_ to 13 mM. Note, to adjust for chelation of Ca^2+^ by aspartate, Low-Cl^-^ HTF contained additional 4.55 mM CaCl_2_, ensuring that the free [Ca^2+^]_o_ was similar to that of HTF ([Ca^2+^]_o_ free = 1.6 mM) as determined by the ABL800 FLEX blood gas analyzer (Radiometer).

To study the action of [Ca^2+^]_o_ on V_rest_, [Ca^2+^]_o_ was adjusted during the final step of the indicator-loading protocol. After washing with standard HTF, the indicator-loaded sperm pellet was resuspended in either 0.5 mM-Ca^2+^ HTF ([Ca^2+^]_o_ free = 0.4 mM), which contained (in mM) 93.8 NaCl, 4.7 KCl, 0.2 MgSO_4_, 0.369 KH_2_PO_4_, 0.5 CaCl_2_, 0.33 Na-pyruvate, 20.98 HEPES, 2.78 glucose, 21.4 Na-lactate, 4 NaHCO_3_, pH 7.35 at 37°C (adjusted with NaOH), or 5 mM-Ca^2+^ HTF ([Ca^2+^]_o_ free = 4 mM), which contained (in mM): 93.8 NaCl, 4.7 KCl, 0.2 MgSO_4_, 0.369 KH_2_PO_4_, 5 CaCl_2_, 0.33 Na-pyruvate, 20.98 HEPES, 2.78 glucose, 21.4 Na-lactate, 4 NaHCO_3_, pH 7.35 at 37°C (adjusted with NaOH). Note, due to chelation of Ca^2+^ by lactate (Vavurusova et al., 2013), the actual free [Ca^2+^]_o_ in these buffers was 0.4 mM and 4 mM, respectively.

To study the action of [Ca^2+^]_o_ on the progesterone-induced V_m_ response, indicator-loaded sperm in standard HTF ([Ca^2+^]_o_ free = 1.6 mM) were mixed in the stopped-flow with either BAPTA-containing Ca^2+^-free HTF or High-Ca^2+^ variants of HTF, resulting in a final free [Ca^2+^]_o_ of 0.15, 0.25, 0.35, 1.6, 4.0, 8.1, 12.2, 16.5, 20.7, or 25.0 mM. To achieve a final free [Ca^2+^]_o_ of < 1.6 mM, sperm were mixed with BAPTA-containing Ca^2+^-free HTF, which contained (in mM): 0.1 – 1.6 BAPTA, 93.8 NaCl, 4.7 KCl, 0.2 MgSO_4_, 0.369 KH_2_PO_4_, 0.33 Na-pyruvate, 20.98 HEPES, 2.78 glucose, 21.4 Na-lactate, 4 NaHCO_3_, pH 7.35 at 30°C (adjusted with NaOH). The free [Ca^2+^]_o_ was confirmed by fluorimetry using Fluo-5N pentapotassium salt (ThermoFisher). To achieve a final free [Ca^2+^]_o_ > 1.6 mM, sperm were mixed with High-Ca^2+^ variants of HTF, which were prepared by mixing standard HTF ([Ca^2+^]_o_ free = 1.6 mM) in varying proportions with High-Ca^2+^ HTF ([Ca^2+^]_o_ free = 53.6 mM), which contained (in mM): 4.7 KCl, 0.2 MgCl_2_, 62 CaCl_2_, 0.33 Na-pyruvate, 22.98 HEPES, 2.78 glucose, 21.4 Na-lactate, 4 NaHCO_3_, pH 7.35 at 30°C (adjusted with NaOH). Note, we report [Ca^2+^]_o_ as the actual free [Ca^2+^]_o_, adjusted for Ca^2+^ chelation by lactate (Vavurusova et al., 2013).

To study low-[Ca^2+^]_o_-induced depolarizations caused by monovalent ion flux via CatSper (Fig. S3), VF2.1.Cl-loaded sperm were mixed in the stopped-flow with BAPTA-containing Ca^2+^-free HTF, which contained in (mM): 1.4 – 3 BAPTA, 93.8 NaCl, 4.7 KCl, 0.2 MgSO_4_, 0.369 KH_2_PO_4_, 0.33 Na-pyruvate, 20.98 HEPES, 2.78 glucose, 21.4 Na-lactate, 4 NaHCO_3_, pH 7.35 at 30°C (adjusted with NaOH). The final free [Ca^2+^]_o_ after mixing was between 0.1 µM and 250 µM, and was confirmed with fluorimetry using Fluo-5N pentapotassium salt (ThermoFisher).

### Spectral acquisition of VF2.1.Cl-loaded human sperm

Excitation and emission spectra of VF2.1.Cl-loaded human sperm (3 x 10^7^ sperm/ml in HTF) were recorded with a fluorescence plate reader (CLARIOStar, BMG) in spectral scanning mode using a bottom optic with a 4 mm focal height in a 384-well plate (Greiner Bio-One) with 50 µl per well.

### Stopped-flow recording of V_m_ and Ca^2+^ signals

Changes in sea urchin sperm V_m_ were measured at 18°C in a stopped-flow device (SFM-4000; Biologic). Indicator-loaded sea urchin sperm were mixed 1:1 (v/v) at a flow rate of 2 ml/s with ASW or [K^+^]_o_-varied ASW supplemented with resact as previously described (Strünker et al., 2006). For experiments with human and boar sperm, changes in V_m_ were measured in a micro stopped-flow device (µSFM, Biologic) at 30°C. Housed in 500 µl glass syringes (Hamilton), the indicator-loaded sperm suspension was mixed 1:1 (v/v) with HTF or ionic variants of HTF supplemented without (baseline) or with valinomycin (4 µM) (Cayman Chemical), progesterone (200 pM – 60 µM) (Cayman Chemical), prostaglandin E1 (4 µM) (Cayman Chemical), NH_4_Cl (60 mM), quinidine (10 µM – 1 mM), or ionomycin (20 µM) (Cayman Chemical). The concentrations of ions and compounds in the results are given as final concentrations after mixing. The mixing flow rate was either 1 ml/s (dead time 1.7 ms) when recording V_m_ signals or 0.5 ml/s (dead time 3.4 ms) when simultaneously recording V_m_ and Ca^2+^ signals. Fluorescence was recorded using FAST^M^ (Kierzek et al., 2021). In brief, fluorescence of indicator-loaded sea urchin sperm was excited by a SpectraX Light Engine (Lumencor) modulated at 10 kHz and fitted with bandpass filters (Table 1). Fluorescence from indicator-loaded human or boar sperm was excited by an array of LEDs (Thorlabs) operated by a custom-made LED driver and modulated by 33.7 kHz. Lock-in amplifiers (MFLI, Zurich Instruments and Model 7230, Signal Recovery) supplied the signals to modulate the LEDs. Modulated excitation light was delivered via a liquid light guide (series 380, Ø3mm x 1000mm, Lumatec) to a cuvette, either FC-15 (Biologic) for sea urchin sperm or µFC-15 (Biologic) for human and boar sperm. The emission was collected at right angles to the cuvette and spectrally filtered with bandpass filters (Table 1) onto two photomultiplier-detection modules (H10723-20; Hamamatsu Photonics). Signals from the PMTs were directed to lock-in amplifiers, where they were amplified and frequency filtered with a 3^rd^ order (18 dB/octave) low pass filter and a time constant of 1 ms. Data acquisition was performed with a data acquisition pad (PCI-6221; National Instruments) and Bio-Kine software (BioLogic) with a sampling rate of 1 or 2 ms. For simultaneous recording of V_m_ and Ca^2+^ signals, VF2.1.Cl was excited with modulated light as described above and Calbryte 630 was excited by light from an LED (M565L3; Thorlabs) fitted with a 586/20 filter (Semrock) modulated by a lock-in amplifier (SR844, Standford Research Systems) at 27.1 kHz. The light intensity, measured with a power meter (PM100USB, Thorlabs), was 10 mW for excitation of VF2.1.Cl and 5.4 mW for excitation of Calbryte 630. Fluorescence was detected at rights angles with PMTs fitted with filters (Table 2) and demodulated by lock-in amplifiers and recorded as described above.

**Table 1.**
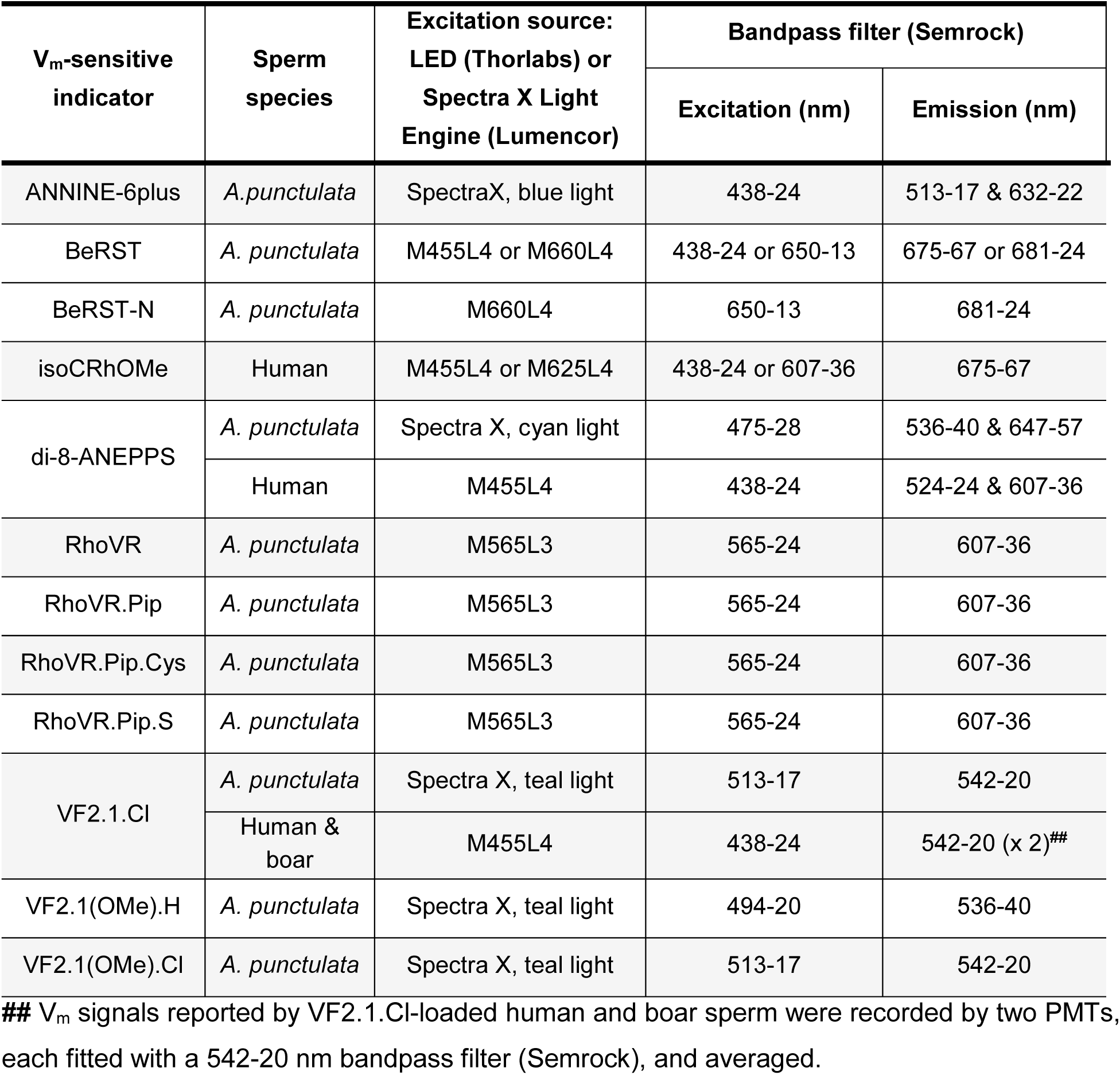
Optical configurations for recording sperm V_m_ signals.

| $V_m$-sensitive indicator | Sperm species | Excitation source:<br>LED (Thorlabs) or<br>Spectra X Light<br>Engine (Lumencor) | Bandpass filter (Semrock) | |
| --- | --- | --- | --- | --- |
|  |  |  | Excitation (nm) | Emission (nm) |
| ANNINE-6plus | *A.punctulata* | SpectraX, blue light | 438-24 | 513-17 & 632-22 |
| BeRST | *A. punctulata* | M455L4 or M660L4 | 438-24 or 650-13 | 675-67 or 681-24 |
| BeRST-N | *A. punctulata* | M660L4 | 650-13 | 681-24 |
| isoCRhOMe | Human | M455L4 or M625L4 | 438-24 or 607-36 | 675-67 |
| di-8-ANEPPS | *A. punctulata* | Spectra X, cyan light | 475-28 | 536-40 & 647-57 |
|  | Human | M455L4 | 438-24 | 524-24 & 607-36 |
| RhoVR | *A. punctulata* | M565L3 | 565-24 | 607-36 |
| RhoVR.Pip | *A. punctulata* | M565L3 | 565-24 | 607-36 |
| RhoVR.Pip.Cys | *A. punctulata* | M565L3 | 565-24 | 607-36 |
| RhoVR.Pip.S | *A. punctulata* | M565L3 | 565-24 | 607-36 |
| VF2.1.Cl | *A. punctulata* | Spectra X, teal light | 513-17 | 542-20 |
|  | Human &<br>boar | M455L4 | 438-24 | 542-20 (x 2)<sup>##</sup> |
| VF2.1(OMe).H | *A. punctulata* | Spectra X, teal light | 494-20 | 536-40 |
| VF2.1(OMe).Cl | *A. punctulata* | Spectra X, teal light | 513-17 | 542-20 |

<sup>##</sup> $V_m$ signals reported by VF2.1.Cl-loaded human and boar sperm were recorded by two PMTs, each fitted with a 542-20 nm bandpass filter (Semrock), and averaged.

**Table 2.** Optical configuration for recording V_m_ and Ca^2+^ signals from human sperm.

| Indicator type | Indicator | Excitation |  | Bandpass filter<br>(Semrock) |  |
| --- | --- | --- | --- | --- | --- |
|  |  | LED<br>(Thorlabs) | Modulation | Excitation<br>(nm) | Emission<br>(nm) |
| $V_m$ | VF2.1.Cl | M455L4 | 33.7 kHz | 438-24 | 542-20 |
| $Ca^{2+}$ | Calbryte 630 | M565L3 | 27.1 kHz | 586/20 | 647-57 |

### Analysis of stopped-flow recorded V_m_ signals

Each V_m_ signal represents the average of at least five recordings. Signals reported by VoltageFluors are depicted as the percent change in fluorescence (ΔF/F_0_ (%)) with respect to the mean of the first 5 - 10 data points after mixing (F_0_); the baseline ΔF/F_0_ obtained upon mixing with ASW (sea urchin sperm) or HTF (human and boar sperm) was subtracted. Signals reported by electrochromic dyes are reported as the percent change in the fluorescence ratio (ΔR/R_0_ (%); R = shorter/longer wavelength) with respect to the mean of the first 5 - 10 data points (R_0_); the baseline ΔR/R_0_ signal was subtracted. The stimulus-induced changes in ΔF/F_0_ (%) or ΔR/R_0_ (%) were rescaled to yield absolute V_m_ values (mV) using the V_rest_ and slope of the fitted line (ΔF/F or ΔR/R % per mV) from an accompanying null-point calibration.

### Null-point calibration for converting fluorescence into millivolts

For sea urchin sperm, the calibration procedure to convert fluorescence into millivolts (mV) has been previously described (Hamzeh et al., 2019; Kierzek et al., 2021; Seifert et al., 2015; Strünker et al., 2006). Briefly, sperm were mixed with [K^+^]_o_-varied ASW supplemented with resact (4 nM). A plot of the peak amplitude (ΔF/F_0_ (%) or ΔR/R_0_ (%)) versus E_K_ for a given [K^+^]_o_ was fit with a linear regression. The slope of the fitted line yielded the percent change in ΔF/F_0_ or ΔR/R_0_ per mV (ΔF/F_0_ or ΔR/R_0_ (%) per 10 mV), and the x-intercept yielded V_rest_. E_K_ was calculated assuming a [K^+^]_i_ of 423 mM (Strünker et al., 2006). For the null-point calibration in human and boar sperm, V_m_ was clamped to different E_K_ values by mixing the sperm with [K^+^]_o_-varied HTF supplemented with the K^+^-ionophore valinomycin (4 µM). A plot of the amplitude (ΔF/F_0_ (%) or ΔR/R_0_ (%)) versus E_K_ was fit with a line. The slope of the line reported the percent change in ΔF/F_0_ or ΔR/R_0_ per mV (ΔF/F (%) or ΔR/R (%) per 10 mV) and the x-intercept revealed the V_rest_. E_K_ was calculated using a [K^+^]_i_ of 120 mM (Fig. S1).

### Dose-response analysis

Progesterone-evoked V_m_ signals were processed and analyzed in Clampfit 11.1 (Molecular Devices) to extract dose-dependent V_m_ signal parameters. The peak V_m_ (V_max_) reached during the depolarization was determined by eye for each dose. The signal amplitude (ΔV_m_) was defined as the difference between V_max_ and V_rest_. To determine the rates of depolarization and repolarization, V_m_ signals were first filtered using a lowpass Gaussian filter with a −3 dB cutoff between 1 - 12 Hz. To determine the depolarization rate, a linear regression was fit to data points residing within 40 – 60% of V_max_. To determine the repolarization rate, the regression was applied to data residing within 70 – 90% of V_max_. For dose-response analysis, the signal amplitude, depolarization rate, and repolarization rate were analyzed using a non-linear regression with a four-parameter fit in GraphPad Prism 9. To generate phase plane plots, the first derivative of the filtered V_m_ signals, reported by capacitated sperm, was determined in Clampfit and plotted against V_m_.

### Analysis of the Ca^2+^-dependence of progesterone-induced voltage responses

To determine the [Ca^2+^]_o_ at which the time-to-peak (the peak V_m_ is also referred to as V_max_) of the progesterone-induced depolarization was half-maximal, a non-linear regression with four-parameter fit was applied using GraphPad Prism 9. To account for the increase in scatter among replicates at low [Ca^2+^]_o_, the fit was applied with a weighting factor of 1/Y^2^. For double logarithmic plots, the logarithm of the mean depolarization rate and the logarithm of the mean repolarization rate were plotted against the logarithm of free [Ca^2+^]_o_ and fit with linear regressions.

### Determination of the relative permeabilities setting the resting membrane potential

The V_rest_ determined at each [K^+^]_o_ was plotted against [K^+^]_o_ and fit with the Goldman-Hodgkin-Katz (GHK) equation. Since V_rest_ was independent of [Cl^-^]_o_ (Fig. 3A), terms for [Cl^-^]_o_ and [Cl^-^]_i_ were omitted from the equation. The GHK equation was fit by setting the K^+^ permeability (P_K_) to 1.0 and solving for the Na^+^ permeability (P_Na_) based on equation 1 below:

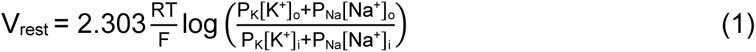

Where, T is the temperature in Kelvins (303.15°K, i.e., 30°C), R is the universal gas constant (8.3145 J·mol^-1^·K^-1^), F is the Faraday constant (96485 C/mol), [K^+^]_i_ is 120 mM and [Na^+^]_i_ is 14 mM (Babcock, 1983). To maintain osmolarity, changes in [K^+^]_o_ required an equimolar substitution of Na^+^ for K^+^, hence [Na^+^]_o_ = 133.5 – [K^+^]_o_. Therefore, the equation above was re-expressed as follows:

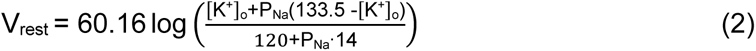

Using GraphPad Prism 9, equation 2 was fit to a plot of the mean V_rest_ against [K^+^]_o_.

### Analysis of Ca^2+^ signals recorded simultaneously with V_m_

Each Ca^2+^ signal represents the average of at least five recordings. Signals are depicted as the percent change in Calbryte 630 fluorescence (ΔF/F_0_ (%)) with respect to the mean of the first 5 - 10 data points after mixing (F_0_); the baseline ΔF/F_0_ obtained upon mixing with HTF was subtracted. The derivative of the Ca^2+^ signal (dCa^2+^/dt) was determined in Clampfit after the Ca^2+^ signal was filtered with a lowpass Gaussian filter with a −3 dB cutoff between 2 - 4 Hz.

### Statistical analysis

Paired comparisons were analyzed using a t-test if the data passed the Shapiro-Wilk normality test; otherwise, a Wilcoxon signed-rank test was used. Unpaired comparisons were analyzed using a two-sample t-test if the data passed the Shapiro-Wilk or Kolmogorov-Smirnov normality test; otherwise, the Mann-Whitney test was used. Asterisks denote statistical significances as follows: * p < 0.05, ** p< 0.01, *** p < 0.001.

## Acknowledgements

We thank present and former members of U. Benjamin Kaupp’s laboratory, especially Luis Alvarez, An Gong, Reinhard Seifert, Hussein Hamzeh, and Fedir Lavryk, for collecting, handling, and shipping sea urchin sperm to support this project. We also thank Albrecht Schwab and his lab for allowing us to use their ABL800 FLEX. C.B., T.S., and U.B.K. are supported by the Deutsche Forschungsgemeinschaft (DFG, German Research Foundation): project numbers 329621271 (CRU326; C.B., T.S.), 5486231 (CRC645 U.B.K., T.S.) and 420653497 (T.S.). T.S. is also supported by the Interdisciplinary Center for Clinical Research Münster (IZKF; Str/014/21) and the Europäischer Fonds für regionale Entwicklung (EFRE; 0400305). E.W.M. is supported by the US National Institutes of Health (R35GM153237), Department of Energy (DE-SC0023184), and the Agilent Biodesign Program. M.K. is supported by Bridging Funding by the Cells in Motion Interfaculty Centre.

## Author contributions

T.S. and C.B conceived the project, M.K. contributed to the conception of the project. T.S., C.B., and M.K. designed the research and interpreted the results. M.K. performed experiments, analyzed the data, and visualized the results. Cr.B., D.F, E.W.M, and U.B.K. contributed to the design of research and/or analysis and interpretation of results. M.K. and T.S. drafted the manuscript. U.B.K. and C.B. contributed to the writing and reviewed and edited the manuscript. All authors revised the manuscript critically for important intellectual content and approved the manuscript.

## Data availability

The data that support the findings of this study are available from the corresponding author upon reasonable request.

## Materials & Correspondence

Correspondence and material requests should be addressed to Timo Strünker

## Supplementary Materials

**Supplementary Figure 1.**
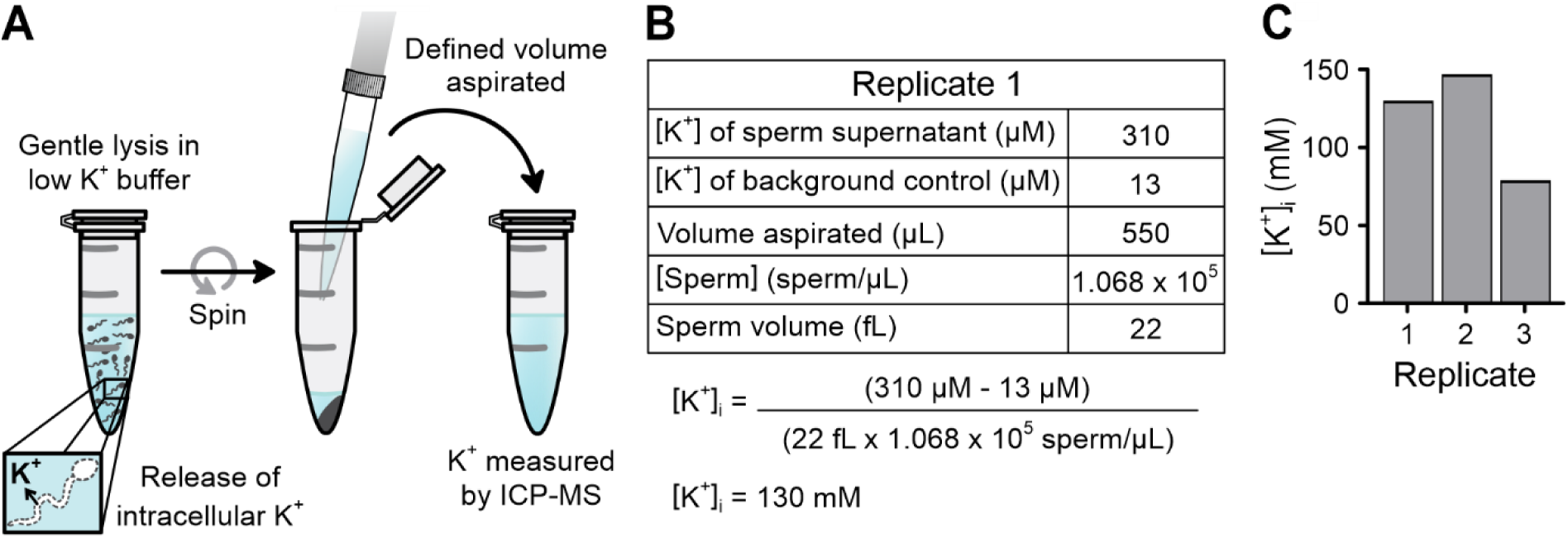
Determination of the intracellular K^+^ concentration in human sperm. **(A)** Schematic depicting how the intracellular K^+^ concentration ([K^+^]_i_) was determined. Intracellular K^+^ from a dense sperm suspension of known concentration was released by permeabilizing the plasma membrane with a hypotonic, low K^+^ buffer containing 0.001% digitonin. This release elevated the K^+^ concentration in the medium. After centrifugation, a defined volume of the supernatant (550 μl) was aspirated, and the K^+^ concentration was determined by inductively coupled mass spectrometry (ICP-MS). **(B)** Results and calculation for one representative replicate. The protocol outlined in (A) was performed in parallel for an equivalent volume of sperm-free medium; this background control accounts for the K^+^ in the medium prior to K^+^ release from sperm. **(C)** Graph showing the [K^+^]_i_ result for each replicate. The mean [K^+^]_i_ was 120 ± 40 mM (n = 3).

**Supplementary Figure 2.**
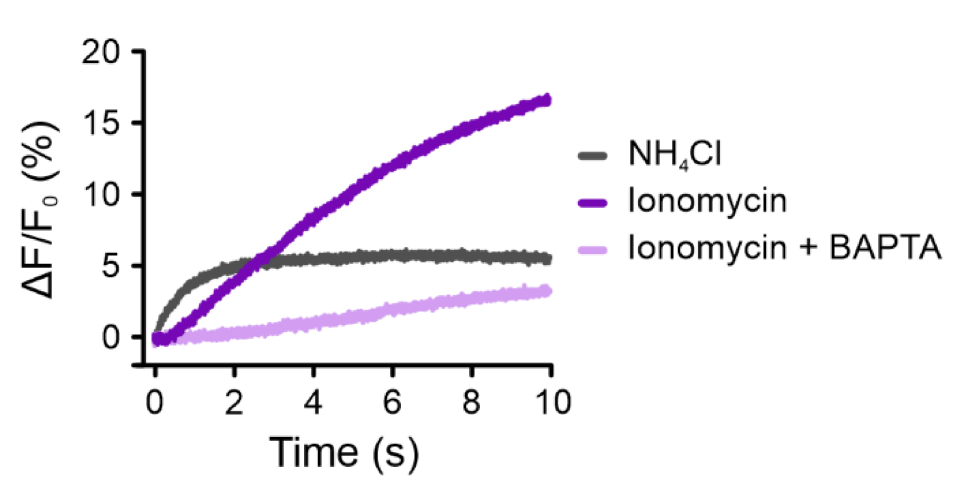
Ionomycin-induced changes in intracellular pH. Increase in intracellular pH (pH_i_) induced by mixing human sperm with ionomycin (10 μM) or NH_4_Cl (30 mM), reported by pHrodo. The pH_i_ response was attenuated when ionomycin was co-delivered with the extracellular Ca^2+^ chelator BAPTA (3.2 mM), adjusting [Ca^2+^]_o_ to < 100 nM. This reinforces that the pH_i_ response arises from the Ca^2+^/H^+^ exchange by ionomycin.

**Supplementary Figure 3.**
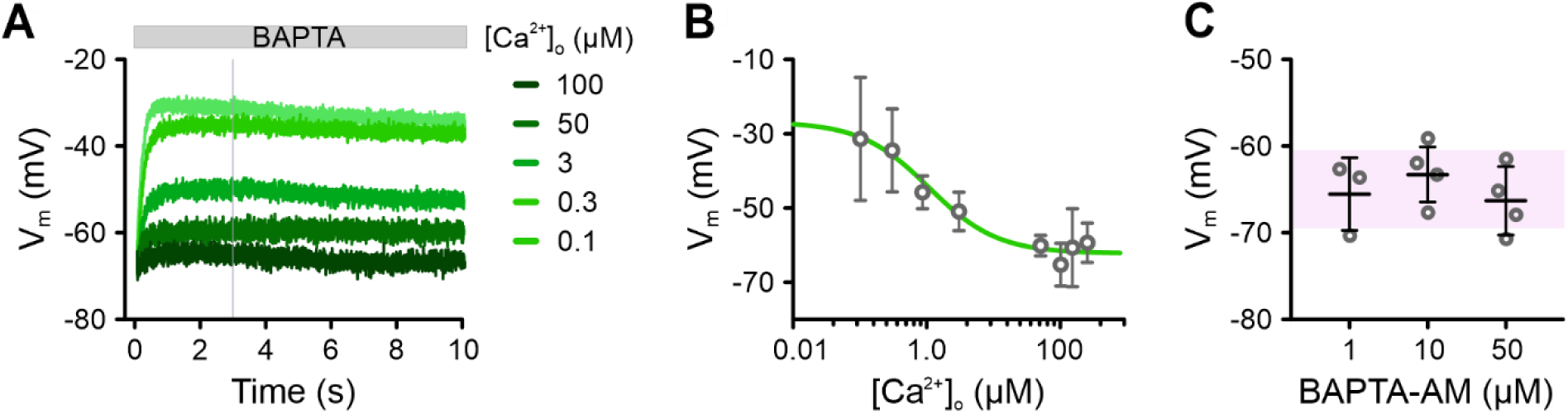
The action of low extracellular Ca^2+^ and buffering of intracellular Ca^2+^ by BAPTA on the membrane potential of human sperm. **(A)** Average (n ≥ 3) changes in V_m_ of non-capacitated VF2.1.Cl-loaded sperm induced by mixing with varying concentrations of BAPTA, reducing extracellular Ca^2+^ ([Ca^2+^]_o_) from 1.6 mM to the values indicated. Error bars were omitted for clarity. Fluorescence was converted into millivolts by accompanying null-point calibrations (not shown). **(B)** Characterization of the relationship between the V_m_ response, averaged over the grey window in (A), and the final [Ca^2+^]_o_. The V_m_ response was attenuated at the half-maximal ‘inhibitory’ concentration (IC_50_) of 1.1 µM [Ca^2+^]_o_, which aligns with the IC_50_ values for the inhibition of monovalent CatSper currents by extracellular Ca^2+^ (Lishko et al., 2011; Smith et al., 2015). **(C)** Mean (± SD) V_rest_ of non-capacitated VF2.1.Cl-loaded sperm after loading for 60 min with either 1 µM, 10 µM, or 50 µM BAPTA-AM. The shaded region shows the V_rest_ (± SD) of control, unloaded sperm from Figure 2F (−65 ± 5 mV; n = 110).

**Supplementary Figure 4.**
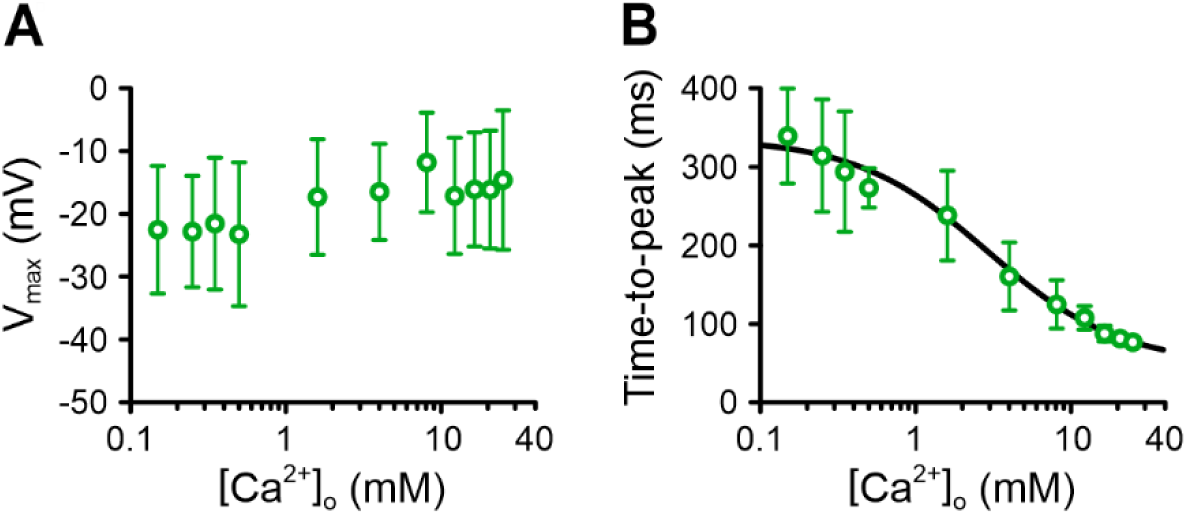
The action of extracellular Ca^2+^ on progesterone-induced V_m_ responses in human sperm. **(A, B)** The mean (± SD) **(A)** V_max_ and **(B)** time-to-peak (peak = V_max_) of progesterone-induced V_m_ responses at varying [Ca^2+^]_o_, shown in Figure 7A and C (n ≥ 4). The time-to-peak was half-maximal (EC_50_) at 3 mM [Ca^2+^]_o_.

**Supplementary Figure 5.**
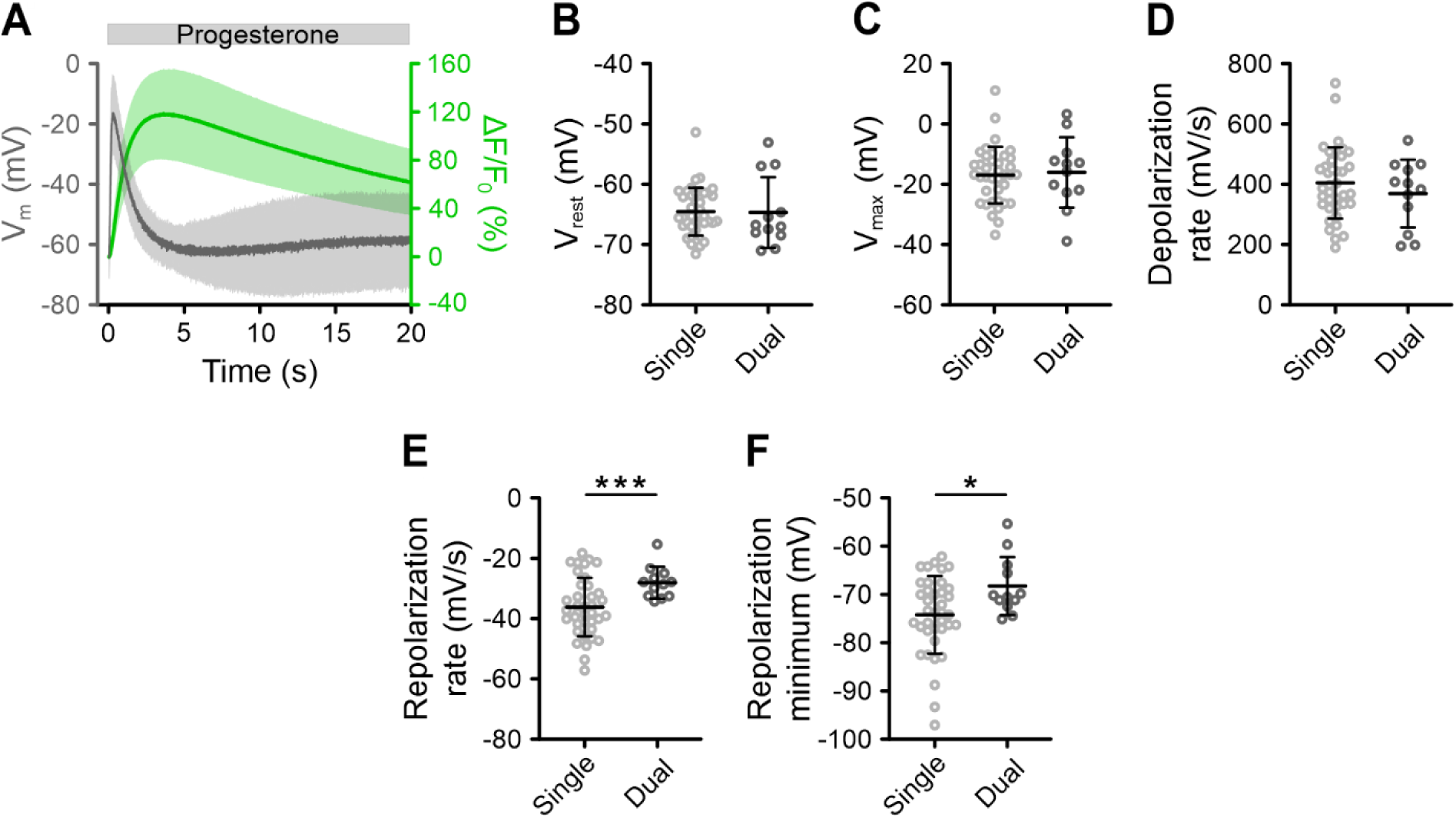
Properties of the progesterone-evoked V_m_ signal recorded individually compared to simultaneously with the Ca^2+^ signal. **(A)** Progesterone-evoked V_m_ and Ca^2+^ signals (mean ± SD; n = 12) in non-capacitated human sperm loaded with VF2.1.Cl and Calbryte 630, recorded simultaneously with FAST^M^ at 1.6 mM [Ca^2+^]_o_. **(B–F)** Mean (± SD) **(B)** V_rest_, **(C)** V_max_, **(D)** depolarization rate, **(E)** repolarization rate, and **(F)** repolarization minimum determined for progesterone-evoked V_m_ signals recorded from VF2.1.Cl-loaded, non-capacitated sperm either with (n = 12) or without (n = 38) loading with the Ca^2+^ indicator Calbryte 630, referred to in the figure as dual and single recording, respectively. Sparse loading with Calbryte 630 resulted in some buffering of [Ca^2+^]_i_, resulting in mild attenuations in the Ca^2+^-dependent repolarization phase. * p < 0.05, *** p < 0.001.

**Supplementary Table 1.** Literature review of previous determinations of non-capacitated and capacitated human sperm V_rest_.

| $V_{rest}$ (mV) | | Methodology | Publications |
| --- | --- | --- | --- |
| non-capacitated | capacitated |  |  |
| -69 ± 2<br>mean ± SD; n = 6 | -36 ± 3<sup>#</sup><br>mean ± SD; n = 4 | Radioactivity assay | Calzada *et al.*, 1988 |
| -75 ± 6<br>mean ± SD; n = 4 | not determined | Radioactivity assay | Calzada & Tellez, 1997 |
| -40 ± 16<br>mean ± SD; n = 7 | not determined | Spectrofluorimetry with 500 nM DiSC<sub>3</sub>(5) | Linares-Hernandez *et al.*, 1998 |
| not determined | -58 ± 2<br>mean ± SEM; n = 12 | Spectrofluorimetry with 250 nM DiSC<sub>3</sub>(5) | Patrat *et al.*, 2002 |
| not determined | -36.5 ± 3.3<br>(mean ± SEM); n = 33 | Patch-clamp under quasi-physiological gradients | Mansell *et al.*, 2014 |
| -17.7 ± 1.8<br>mean ± SEM; n = 10 | -22.7 ± 2<br>mean ± SEM; n = 16 | Patch-clamp under quasi-physiological gradients | Brown *et al.*, 2016 |
| -63.5<br>n = 1 | not determined | Spectrofluorimetry with 1 µM DiSC<sub>3</sub>(5) | Baro Graf *et al.*, 2019 |
| -63.1<br>n = 1 | -74.5<br>n = 1 | Flow cytometry with 50 nM DiSC<sub>3</sub>(5) |  |
| -37.7 ± 9.9<br>mean ± SD; n = 29 | -57.8 ± 12.9<br>mean ± SD; n = 29 | Spectrofluorimetry with 1 µM DiSC<sub>3</sub>(5) | Baro Graf *et al.*, 2020 |
| -55 ± 14<sup>#</sup><br>mean ± SD; n = 18 | -38 ± 5<sup>#</sup><br>mean ± SD; n = 18 |  |  |
| -39.7 ± 8<sup>#</sup><br>mean ± SD; n = 13 | -38.9 ± 10<sup>#</sup><br>mean ± SD; n = 13 |  |  |
| -35.7 ± 2.8<br>mean ± SEM; n = 13 | -45.2 ± 3.2<br>mean ± SEM; n = 13 | Flow cytometry with 5 nM DiSC<sub>3</sub>(5) | Molina *et al.*, 2020 |
| -67 ± 4<br>mean ± SEM; n = 9 | -70 ± 3<br>mean ± SEM; n = 9 | Flow cytometry with 25 nM DiSC<sub>3</sub>(5) | Matamoros-Volante *et al.*, 2020 |
| -60 ± 3<br>mean ± SEM; n = 6 | -71 ± 3<br>mean ± SEM; n = 6 |  |  |

Publications are listed chronologically and colour-coded based on method. Green: radioactivity assay, yellow: spectrofluorimetry with DiSC<sub>3</sub>(5), orange: flow cytometry with DiSC<sub>3</sub>(5), and blue: sperm patch-clamp under quasi-physiological conditions.

**Supplementary Table 2.** Literature review of non-capacitated and capacitated mouse sperm V_rest_.

| $V_{rest}$ (mV) | | Methodology | Publication |
| --- | --- | --- | --- |
| non-capacitated | capacitated |  |  |
| -42 ± 8.8<br>mean ± SD; n = 16 | not determined | Spectrofluorimetry with 1 µM DiSC<sub>3</sub>(5) | Espinosa & Darszon, 1995 |
| -38 ± 4<br>mean ± SD; n = 3 | -55 ± 2<br>mean ± SD; n = 4 | Spectrofluorimetry with 250 nM DiSC<sub>3</sub>(5) | Zeng *et al.*, 1995 |
| -33 ± 4<br>mean ± SD; n = 3 | -49 ± 3<br>mean ± SD; n = 3 | Spectrofluorimetry with 500 nM DiSBAC<sub>2</sub>(3) |  |
| -25.7 ± 6.6<br>mean ± SD; n = 75 | -63.4 ± 17.1<br>mean ± SD; n = 61 | Single cell fluorimetry with 1 µM di-8-ANEPPS | Arnoult *et al.*, 1999 |
| -52 ± 6<br>mean ± SD; n = 10 | -66 ± 9 mV<br>mean ± SD; n = 10 | Spectrofluorimetry with 1 µM DiSC<sub>3</sub>(5) | Munoz-Garay *et al.*, 2001 |
| -36 ± 2<sup>#</sup><br>mean ± SEM; n = 3 | -67 ± 2<sup>#</sup><br>mean ± SEM; n = 3 | Spectrofluorimetry with 1 µM DiSC<sub>3</sub>(5) | Demarco *et al.*, 2003 |
| -47 ± 3.3 mV<br>mean ± SEM; n = 4 | -62.2 ± 2.9<br>mean ± SEM; n = 4 | Spectrofluorimetry with 1 µM DiSC<sub>3</sub>(5) | Santi *et al.*, 2010 |
| -45.59 ± 2.57<br>mean ± SEM; n = 11 | -63.88 ± 2.92<br>mean ± SEM; n = 11 | Spectrofluorimetry with 1 µM DiSC<sub>3</sub>(5) | Chavez *et al.*, 2013 |
| -42.6<br>n = 1 | -62.4<br>n = 1 | Spectrofluorimetry with 1 µM DiSC<sub>3</sub>(5) | Escoffier *et al.*, 2015 |
| not determined | -59.4 ± 5.3<br>mean ± SEM; n = ? | Flow cytometry with 5 nM DiSC<sub>3</sub>(5) | Molina *et al.*, 2020 |

Publications are listed chronologically and colour-coded according to method: Yellow: spectrofluorimetry with DiSC<sub>3</sub>(5), orange: flow cytometry with DiSC<sub>3</sub>(5), pink: single-cell fluorimetry with di-8-ANEPPS, grey: spectrofluorimetry with DiSBAC<sub>2</sub>(3). Although mouse sperm $V_m$ has been extensively studied by sperm patch-clamp, these publications were excluded from the table as recordings were not performed under physiological conditions at pH 6.5.
<sup>#</sup> Indicates that values were estimated from figures as they were not explicitly given.
? Indicates that n values could not be identified.

## References

Arnoult C, Kazam IG, Visconti PE, Kopf GS, Villaz M, Florman HM. Control of the low voltage-activated calcium channel of mouse sperm by egg ZP3 and by membrane hyperpolarization during capacitation. Proc Natl Acad Sci U S A. 1999 Jun 8;96(12):6757–62. doi: 10.1073/pnas.96.12.6757. PMID: 10359785; PMCID: PMC21988.

Baro Graf C, Ritagliati C, Stival C, Balestrini PA, Buffone MG, Krapf D. Determination of a Robust Assay for Human Sperm Membrane Potential Analysis. Front Cell Dev Biol. 2019 Jun 11;7:101. doi: 10.3389/fcell.2019.00101. PMID: 31245370; PMCID: PMC6579818.

Baro Graf C, Ritagliati C, Torres-Monserrat V, Stival C, Carizza C, Buffone MG, Krapf D. Membrane Potential Assessment by Fluorimetry as a Predictor Tool of Human Sperm Fertilizing Capacity. Front Cell Dev Biol. 2020 Jan 17;7:383. doi: 10.3389/fcell.2019.00383. PMID: 32010695; PMCID: PMC6979052.

Berkefeld H, Sailer CA, Bildl W, Rohde V, Thumfart JO, Eble S, Klugbauer N, Reisinger E, Bischofberger J, Oliver D, Knaus HG, Schulte U, Fakler B. BKCa-Cav channel complexes mediate rapid and localized Ca2+-activated K+ signaling. Science. 2006 Oct 27;314(5799):615–20. doi: 10.1126/science.1132915. PMID: 17068255.

Blackmore PF, Beebe SJ, Danforth DR, Alexander N. Progesterone and 17 alpha-hydroxyprogesterone. Novel stimulators of calcium influx in human sperm. J Biol Chem. 1990 Jan 25;265(3):1376–80. PMID: 2104840.

Bönigk W, Loogen A, Seifert R, Kashikar N, Klemm C, Krause E, Hagen V, Kremmer E, Strünker T, Kaupp UB. An atypical CNG channel activated by a single cGMP molecule controls sperm chemotaxis. Sci Signal. 2009 Oct 27;2(94):ra68. doi: 10.1126/scisignal.2000516. PMID: 19861689.

Brenker C, Goodwin N, Weyand I, Kashikar ND, Naruse M, Krähling M, Müller A, Kaupp UB, Strünker T. The CatSper channel: a polymodal chemosensor in human sperm. EMBO J. 2012 Apr 4;31(7):1654–65. doi: 10.1038/emboj.2012.30. Epub 2012 Feb 21. PMID: 22354039; PMCID: PMC3321208.

Brenker C, Zhou Y, Müller A, Echeverry FA, Trötschel C, Poetsch A, Xia XM, Bönigk W, Lingle CJ, Kaupp UB, Strünker T. The Ca^2+^-activated K^+^ current of human sperm is mediated by Slo3. Elife. 2014 Mar 26;3:e01438. doi: 10.7554/eLife.01438. PMID: 24670955; PMCID: PMC3966514.

Brown SG, Publicover SJ, Mansell SA, Lishko PV, Williams HL, Ramalingam M, Wilson SM, Barratt CL, Sutton KA, Da Silva SM. Depolarization of sperm membrane potential is a common feature of men with subfertility and is associated with low fertilization rate at IVF. Hum Reprod. 2016 Jun;31(6):1147–57. doi: 10.1093/humrep/dew056. Epub 2016 Apr 6. PMID: 27052499; PMCID: PMC4871192.

Brown SG, Publicover SJ, Barratt CLR, Martins da Silva SJ. Human sperm ion channel (dys)function: implications for fertilization. Hum Reprod Update. 2019 Nov 5;25(6):758–776. doi: 10.1093/humupd/dmz032. PMID: 31665287; PMCID: PMC6847974.

Budde T, Meuth S, Pape HC. Calcium-dependent inactivation of neuronal calcium channels. Nat Rev Neurosci. 2002 Nov;3(11):873–83. doi: 10.1038/nrn959. PMID: 12415295.

Calzada L, Bernal A, Loustaunau E. Effect of steroid hormones and capacitation on membrane potential of human spermatozoa. Arch Androl. 1988;21(2):121–8. doi: 10.3109/01485018808986722. PMID: 3223786.

Calzada L, Tellez J. Defective function of membrane potential (psi) on sperm of infertile men. Arch Androl. 1997 Mar-Apr;38(2):151-5. doi: 10.3109/01485019708987892. PMID: 9049036.

Chávez JC, de la Vega-Beltrán JL, Escoffier J, Visconti PE, Treviño CL, Darszon A, Salkoff L, Santi CM. Ion permeabilities in mouse sperm reveal an external trigger for SLO3-dependent hyperpolarization. PLoS One. 2013;8(4):e60578. doi: 10.1371/journal.pone.0060578. Epub 2013 Apr 5. PMID: 23577126; PMCID: PMC3618424.

Chung JJ, Shim SH, Everley RA, Gygi SP, Zhuang X, Clapham DE. Structurally distinct Ca(2+) signaling domains of sperm flagella orchestrate tyrosine phosphorylation and motility. Cell. 2014 May 8;157(4):808–22. doi: 10.1016/j.cell.2014.02.056. PMID: 24813608; PMCID: PMC4032590.

Chung JJ, Miki K, Kim D, Shim SH, Shi HF, Hwang JY, Cai X, Iseri Y, Zhuang X, Clapham DE. CatSperζ regulates the structural continuity of sperm Ca^2+^ signaling domains and is required for normal fertility. Elife. 2017 Feb 23;6:e23082. doi: 10.7554/eLife.23082. PMID: 28226241; PMCID: PMC5362262.

Contet C, Goulding SP, Kuljis DA, Barth AL. BK Channels in the Central Nervous System. Int Rev Neurobiol. 2016;128:281–342. doi: 10.1016/bs.irn.2016.04.001. Epub 2016 May 13. PMID: 27238267; PMCID: PMC4902275.

Curry MR, Millar JD, Tamuli SM, Watson PF. Surface area and volume measurements for ram and human spermatozoa. Biol Reprod. 1996 Dec;55(6):1325–32. doi: 10.1095/biolreprod55.6.1325. PMID: 8949890.

Deal PE, Kulkarni RU, Al-Abdullatif SH, Miller EW. Isomerically Pure Tetramethylrhodamine Voltage Reporters. J Am Chem Soc. 2016 Jul 27;138(29):9085–8. doi: 10.1021/jacs.6b05672. Epub 2016 Jul 18. PMID: 27428174; PMCID: PMC5222532.

Deal PE, Liu P, Al-Abdullatif SH, Muller VR, Shamardani K, Adesnik H, Miller EW. Covalently Tethered Rhodamine Voltage Reporters for High Speed Functional Imaging in Brain Tissue. J Am Chem Soc. 2020 Jan 8;142(1):614–622. doi: 10.1021/jacs.9b12265. Epub 2019 Dec 26. PMID: 31829585; PMCID: PMC6949409.

Ded L, Hwang JY, Miki K, Shi HF, Chung JJ. 3D in situ imaging of the female reproductive tract reveals molecular signatures of fertilizing spermatozoa in mice. Elife. 2020 Oct 20;9:e62043. doi: 10.7554/eLife.62043. PMID: 33078708; PMCID: PMC7707823.

De La Vega-Beltran JL, Sánchez-Cárdenas C, Krapf D, Hernandez-González EO, Wertheimer E, Treviño CL, Visconti PE, Darszon A. Mouse sperm membrane potential hyperpolarization is necessary and sufficient to prepare sperm for the acrosome reaction. J Biol Chem. 2012 Dec 28;287(53):44384–93. doi: 10.1074/jbc.M112.393488. Epub 2012 Oct 24. PMID: 23095755; PMCID: PMC3531752.

Demarco IA, Espinosa F, Edwards J, Sosnik J, De La Vega-Beltran JL, Hockensmith JW, Kopf GS, Darszon A, Visconti PE. Involvement of a Na^+/HCO3-^ cotransporter in mouse sperm capacitation. J Biol Chem. 2003 Feb 28;278(9):7001–9. doi: 10.1074/jbc.M206284200. Epub 2002 Dec 19. PMID: 12496293.

Erdahl WL, Chapman CJ, Taylor RW, Pfeiffer DR. Ca^2+^ transport properties of ionophores A23187, ionomycin, and 4-BrA23187 in a well defined model system. Biophys J. 1994 May;66(5):1678–93. doi: 10.1016/S0006-3495(94)80959-2. PMID: 8061216; PMCID: PMC1275887.

Escoffier J, Navarrete F, Haddad D, Santi CM, Darszon A, Visconti PE. Flow cytometry analysis reveals that only a subpopulation of mouse sperm undergoes hyperpolarization during capacitation. Biol Reprod. 2015 May;92(5):121. doi: 10.1095/biolreprod.114.127266. Epub 2015 Apr 8. PMID: 25855261; PMCID: PMC4645980.

Espinosa F, Darszon A. Mouse sperm membrane potential: changes induced by Ca^2+^. FEBS Lett. 1995 Sep 18;372(1):119–25. doi: 10.1016/0014-5793(95)00962-9. PMID: 7556631.

Fromherz P, Hübener G, Kuhn B, Hinner MJ. ANNINE-6plus, a voltage-sensitive dye with good solubility, strong membrane binding and high sensitivity. Eur Biophys J. 2008 Apr;37(4):509–14. doi: 10.1007/s00249-007-0210-y. Epub 2007 Aug 9. PMID: 17687549; PMCID: PMC2755735.

Garcia MA, Meizel S. Determination of the steady-state intracellular chloride concentration in capacitated human spermatozoa. J Androl. 1999 Jan-Feb;20(1):88-93. Erratum in: J Androl 1999 May-Jun;20(3):434. PMID: 10100478.

Geng Y, Ferreira JJ, Dzikunu V, Butler A, Lybaert P, Yuan P, Magleby KL, Salkoff L, Santi CM. A genetic variant of the sperm-specific SLO3 K^+^ channel has altered pH and Ca^2+^ sensitivities. J Biol Chem. 2017 May 26;292(21):8978–8987. doi: 10.1074/jbc.M117.776013. Epub 2017 Apr 4. PMID: 28377504; PMCID: PMC5448129.

Gest AMM, Lazzari-Dean JR, Ortiz G, Yaeger-Weiss SK, Boggess SC, Miller EW. A red-emitting carborhodamine for monitoring and measuring membrane potential. Proc Natl Acad Sci U S A. 2024 Apr 2;121(14):e2315264121. doi: 10.1073/pnas.2315264121. Epub 2024 Mar 29. PMID: 38551837; PMCID: PMC10998576.

Hamzeh H, Alvarez L, Strünker T, Kierzek M, Brenker C, Deal PE, Miller EW, Seifert R, Kaupp UB. Kinetic and photonic techniques to study chemotactic signaling in sea urchin sperm. Methods Cell Biol. 2019;151:487–517. doi: 10.1016/bs.mcb.2018.12.001. Epub 2019 Mar 4. PMID: 30948028.

Harper CV, Kirkman-Brown JC, Barratt CL, Publicover SJ. Encoding of progesterone stimulus intensity by intracellular [Ca^2+^] ([Ca^2+^]_i_) in human spermatozoa. Biochem J. 2003 Jun 1;372(Pt 2):407–17. doi: 10.1042/BJ20021560. PMID: 12614198; PMCID: PMC1223411.

Herrmann L, Brenker C, Mittermair T, Bojovic V, Münchow J, Zhu WF, Trugge C, Fußhöller D, Jikeli J, Temme L, Kaupp UB, Strünker T. Chemosensory signalling in human sperm is controlled by Ca^2+^ influx via CatSper and Ca^2+^ clearance via plasma membrane Ca^2+^ ATPases. Br J Pharmacol. 2025 Jun;182(12):2694–2712. doi: 10.1111/bph.70009. Epub 2025 Feb 27. PMID: 40016153.

Hladky SB, Rink TJ. Potential difference and the distribution of ions across the human red blood cell membrane; a study of the mechanism by which the fluorescent cation, diS-C3-(5) reports membrane potential. J Physiol. 1976 Dec;263(2):287-319. doi: 10.1113/jphysiol.1976.sp011632. PMID: 14255; PMCID: PMC1307701.

Huang YL, Walker AS, Miller EW. A Photostable Silicon Rhodamine Platform for Optical Voltage Sensing. J Am Chem Soc. 2015 Aug 26;137(33):10767–76. doi: 10.1021/jacs.5b06644. Epub 2015 Aug 13. PMID: 26237573; PMCID: PMC4666802.

Hwang JY, Mannowetz N, Zhang Y, Everley RA, Gygi SP, Bewersdorf J, Lishko PV, Chung JJ. Dual Sensing of Physiologic pH and Calcium by EFCAB9 Regulates Sperm Motility. Cell. 2019 May 30;177(6):1480–1494.e19. doi: 10.1016/j.cell.2019.03.047. Epub 2019 May 2. PMID: 31056283; PMCID: PMC8808721.

Hwang JY, Chung JJ. CatSper Calcium Channels: 20 Years On. Physiology (Bethesda). 2023 May 1;38(3):0. doi: 10.1152/physiol.00028.2022. Epub 2022 Dec 13. PMID: 36512352; PMCID: PMC10085559.

Jeschke JK, Biagioni C, Schierling T, Wagner IV, Börgel F, Schepmann D, Schüring A, Kulle AE, Holterhus PM, von Wolff M, Wünsch B, Nordhoff V, Strünker T, Brenker C. The Action of Reproductive Fluids and Contained Steroids, Prostaglandins, and Zn^2+^ on CatSper Ca^2+^ Channels in Human Sperm. Front Cell Dev Biol. 2021 Jul 26;9:699554. doi: 10.3389/fcell.2021.699554. PMID: 34381781; PMCID: PMC8350739.

Kaupp UB, Strünker T. Signaling in Sperm: More Different than Similar. Trends Cell Biol. 2017 Feb;27(2):101–109. doi: 10.1016/j.tcb.2016.10.002. Epub 2016 Nov 5. PMID: 27825709.

Kazemipour A, Novak O, Flickinger D, Marvin JS, Abdelfattah AS, King J, Borden PM, Kim JJ, Al-Abdullatif SH, Deal PE, Miller EW, Schreiter ER, Druckmann S, Svoboda K, Looger LL, Podgorski K. Kilohertz frame-rate two-photon tomography. Nat Methods. 2019 Aug;16(8):778–786. doi: 10.1038/s41592-019-0493-9. Epub 2019 Jul 29. Erratum in: Nat Methods. 2019 Sep;16(9):932. doi: 10.1038/s41592-019-0545-1. PMID: 31363222; PMCID: PMC6754705.

Kierzek M, Deal PE, Miller EW, Mukherjee S, Wachten D, Baumann A, Kaupp UB, Strünker T, Brenker C. Simultaneous recording of multiple cellular signaling events by frequency- and spectrally-tuned multiplexing of fluorescent probes. Elife. 2021 Dec 3;10:e63129. doi: 10.7554/eLife.63129. PMID: 34859780; PMCID: PMC8700268.

Kirkman-Brown JC, Bray C, Stewart PM, Barratt CL, Publicover SJ. Biphasic elevation of [Ca^(2+)^]_(i)_ in individual human spermatozoa exposed to progesterone. Dev Biol. 2000 Jun 15;222(2):326–35. doi: 10.1006/dbio.2000.9729. PMID: 10837122.

Linares-Hernández L, Guzmán-Grenfell AM, Hicks-Gomez JJ, González-Martínez MT. Voltage-dependent calcium influx in human sperm assessed by simultaneous optical detection of intracellular calcium and membrane potential. Biochim Biophys Acta. 1998 Jun 24;1372(1):1–12. doi: 10.1016/s0005-2736(98)00035-2. PMID: 9651467.

Lishko PV, Botchkina IL, Kirichok Y. Progesterone activates the principal Ca2+ channel of human sperm. Nature. 2011 Mar 17;471(7338):387–91. doi: 10.1038/nature09767. PMID: 21412339.

Liu P, Miller EW. Electrophysiology, Unplugged: Imaging Membrane Potential with Fluorescent Indicators. Acc Chem Res. 2020 Jan 21;53(1):11–19. doi: 10.1021/acs.accounts.9b00514. Epub 2019 Dec 13. PMID: 31834772; PMCID: PMC7266091.

Liu B, Mundt N, Miller M, Clapham DE, Kirichok Y, Lishko PV. Recording Electrical Currents across the Plasma Membrane of Mammalian Sperm Cells. J Vis Exp. 2021 Feb 14;(168). doi: 10.3791/62049. PMID: 33645583.

Loew LM. Design and Use of Organic Voltage Sensitive Dyes. Adv Exp Med Biol. 2015;859:27–53. doi: 10.1007/978-3-319-17641-3_2. PMID: 26238048.

Luo T, Chen HY, Zou QX, Wang T, Cheng YM, Wang HF, Wang F, Jin ZL, Chen Y, Weng SQ, Zeng XH. A novel copy number variation in CATSPER2 causes idiopathic male infertility with normal semen parameters. Hum Reprod. 2019 Mar 1;34(3):414–423. doi: 10.1093/humrep/dey377. PMID: 30629171.

Luque GM, Schiavi-Ehrenhaus LJ, Jabloñski M, Balestrini PA, Novero AG, Torres NI, Osycka-Salut CE, Darszon A, Krapf D, Buffone MG. High-throughput screening method for discovering CatSper inhibitors using membrane depolarization caused by external calcium chelation and fluorescent cell barcoding. Front Cell Dev Biol. 2023 Jan 19;11:1010306. doi: 10.3389/fcell.2023.1010306. PMID: 36743410; PMCID: PMC9892719.

Lv M, Liu C, Ma C, Yu H, Shao Z, Gao Y, Liu Y, Wu H, Tang D, Tan Q, Zhang J, Li K, Xu C, Geng H, Zhang J, Li H, Mao X, Ge L, Fu F, Zhong K, Xu Y, Tao F, Zhou P, Wei Z, He X, Zhang F, Cao Y. Homozygous mutation in SLO3 leads to severe asthenoteratozoospermia due to acrosome hypoplasia and mitochondrial sheath malformations. Reprod Biol Endocrinol. 2022 Jan 3;20(1):5. doi: 10.1186/s12958-021-00880-4. PMID: 34980136; PMCID: PMC8722334.

Lyon M, Li P, Ferreira JJ, Lazarenko RM, Kharade SV, Kramer M, McClenahan SJ, Days E, Bauer JA, Spitznagel BD, Weaver CD, Borrego Alvarez A, Puga Molina LC, Lybaert P, Khambekar S, Liu A, Lindsley CW, Denton J, Santi CM. A selective inhibitor of the sperm-specific potassium channel SLO3 impairs human sperm function. Proc Natl Acad Sci U S A. 2023 Jan 24;120(4):e2212338120. doi: 10.1073/pnas.2212338120. Epub 2023 Jan 17. PMID: 36649421; PMCID: PMC9942793.

Mansell SA, Publicover SJ, Barratt CL, Wilson SM. Patch clamp studies of human sperm under physiological ionic conditions reveal three functionally and pharmacologically distinct cation channels. Mol Hum Reprod. 2014 May;20(5):392–408. doi: 10.1093/molehr/gau003. Epub 2014 Jan 16. PMID: 24442342; PMCID: PMC4004083.

Martin G, Sabido O, Durand P, Levy R. Phosphatidylserine externalization in human sperm induced by calcium ionophore A23187: relationship with apoptosis, membrane scrambling and the acrosome reaction. Hum Reprod. 2005 Dec;20(12):3459–68. doi: 10.1093/humrep/dei245. Epub 2005 Aug 19. PMID: 16113043.

Matamoros-Volante A, Castillo-Viveros V, Torres-Rodríguez P, Treviño MB, Treviño CL. Time-Lapse Flow Cytometry: A Robust Tool to Assess Physiological Parameters Related to the Fertilizing Capability of Human Sperm. Int J Mol Sci. 2020 Dec 24;22(1):93. doi: 10.3390/ijms22010093. PMID: 33374265; PMCID: PMC7796328.

Miller EW, Lin JY, Frady EP, Steinbach PA, Kristan WB Jr, Tsien RY. Optically monitoring voltage in neurons by photo-induced electron transfer through molecular wires. Proc Natl Acad Sci U S A. 2012 Feb 7;109(6):2114–9. doi: 10.1073/pnas.1120694109. Epub 2012 Jan 24. PMID: 22308458; PMCID: PMC3277584.

Miyata H, Satouh Y, Mashiko D, Muto M, Nozawa K, Shiba K, Fujihara Y, Isotani A, Inaba K, Ikawa M. Sperm calcineurin inhibition prevents mouse fertility with implications for male contraceptive. Science. 2015 Oct 23;350(6259):442–5. doi: 10.1126/science.aad0836. Epub 2015 Oct 1. PMID: 26429887.

Molina LC, Gunderson S, Riley J, Lybaert P, Borrego-Alvarez A, Jungheim ES, Santi CM. Membrane Potential Determined by Flow Cytometry Predicts Fertilizing Ability of Human Sperm. Front Cell Dev Biol. 2020 Jan 21;7:387. doi: 10.3389/fcell.2019.00387. PMID: 32039203; PMCID: PMC6985285.

Muñoz-Garay C, De la Vega-Beltrán JL, Delgado R, Labarca P, Felix R, Darszon A. Inwardly rectifying K(+) channels in spermatogenic cells: functional expression and implication in sperm capacitation. Dev Biol. 2001 Jun 1;234(1):261–74. doi: 10.1006/dbio.2001.0196. PMID: 11356034.

Okunade GW, Miller ML, Pyne GJ, Sutliff RL, O’Connor KT, Neumann JC, Andringa A, Miller DA, Prasad V, Doetschman T, Paul RJ, Shull GE. Targeted ablation of plasma membrane Ca2+-ATPase (PMCA) 1 and 4 indicates a major housekeeping function for PMCA1 and a critical role in hyperactivated sperm motility and male fertility for PMCA4. J Biol Chem. 2004 Aug 6;279(32):33742–50. doi: 10.1074/jbc.M404628200. Epub 2004 Jun 3. PMID: 15178683.

Patrat C, Serres C, Jouannet P. Progesterone induces hyperpolarization after a transient depolarization phase in human spermatozoa. Biol Reprod. 2002 Jun;66(6):1775–80. doi: 10.1095/biolreprod66.6.1775. PMID: 12021061.

Plásek J, Hrouda V. Assessment of membrane potential changes using the carbocyanine dye, diS-C3-(5): synchronous excitation spectroscopy studies. Eur Biophys J. 1991;19(4):183-8. doi: 10.1007/BF00196344. PMID: 2029874.

Rahban R, Nef S. CatSper: The complex main gate of calcium entry in mammalian spermatozoa. Mol Cell Endocrinol. 2020 Dec 1;518:110951. doi: 10.1016/j.mce.2020.110951. Epub 2020 Jul 24. PMID: 32712386.

Rehfeld A, Mendoza N, Ausejo R, Skakkebæk NE. Bisphenol A Diglycidyl Ether (BADGE) and Progesterone Do Not Induce Ca^2+^ Signals in Boar Sperm Cells. Front Physiol. 2020 Jul 7;11:785. doi: 10.3389/fphys.2020.00785. PMID: 32774306; PMCID: PMC7381341.

Ritagliati C, Baro Graf C, Stival C, Krapf D. Regulation mechanisms and implications of sperm membrane hyperpolarization. Mech Dev. 2018 Dec;154:33–43. doi: 10.1016/j.mod.2018.04.004. Epub 2018 Apr 22. PMID: 29694849.

Santi CM, Martínez-López P, de la Vega-Beltrán JL, Butler A, Alisio A, Darszon A, Salkoff L. The SLO3 sperm-specific potassium channel plays a vital role in male fertility. FEBS Lett. 2010 Mar 5;584(5):1041–6. doi: 10.1016/j.febslet.2010.02.005. Epub 2010 Feb 9. PMID: 20138882; PMCID: PMC2875124.

Schiffer C, Müller A, Egeberg DL, Alvarez L, Brenker C, Rehfeld A, Frederiksen H, Wäschle B, Kaupp UB, Balbach M, Wachten D, Skakkebaek NE, Almstrup K, Strünker T. Direct action of endocrine disrupting chemicals on human sperm. EMBO Rep. 2014 Jul;15(7):758–65. doi: 10.15252/embr.201438869. Epub 2014 May 12. PMID: 24820036; PMCID: PMC4196979.

Schiffer C, Rieger S, Brenker C, Young S, Hamzeh H, Wachten D, Tüttelmann F, Röpke A, Kaupp UB, Wang T, Wagner A, Krallmann C, Kliesch S, Fallnich C, Strünker T. Rotational motion and rheotaxis of human sperm do not require functional CatSper channels and transmembrane Ca^2+^ signaling. EMBO J. 2020 Feb 17;39(4):e102363. doi: 10.15252/embj.2019102363. Epub 2020 Jan 19. PMID: 31957048; PMCID: PMC7024840.

Schuh K, Cartwright EJ, Jankevics E, Bundschu K, Liebermann J, Williams JC, Armesilla AL, Emerson M, Oceandy D, Knobeloch KP, Neyses L. Plasma membrane Ca^2+^ ATPase 4 is required for sperm motility and male fertility. J Biol Chem. 2004 Jul 2;279(27):28220–6. doi: 10.1074/jbc.M312599200. Epub 2004 Apr 12. PMID: 15078889.

Seifert R, Flick M, Bönigk W, Alvarez L, Trötschel C, Poetsch A, Müller A, Goodwin N, Pelzer P, Kashikar ND, Kremmer E, Jikeli J, Timmermann B, Kuhl H, Fridman D, Windler F, Kaupp UB, Strünker T. The CatSper channel controls chemosensation in sea urchin sperm. EMBO J. 2015 Feb 3;34(3):379–92. doi: 10.15252/embj.201489376. Epub 2014 Dec 22. PMID: 25535245; PMCID: PMC4339123.

Smith JF, Syritsyna O, Fellous M, Serres C, Mannowetz N, Kirichok Y, Lishko PV. Disruption of the principal, progesterone-activated sperm Ca^2+^ channel in a CatSper2-deficient infertile patient. Proc Natl Acad Sci U S A. 2013 Apr 23;110(17):6823–8. doi: 10.1073/pnas.1216588110. Epub 2013 Mar 25. PMID: 23530196; PMCID: PMC3637729.

Strünker T, Weyand I, Bönigk W, Van Q, Loogen A, Brown JE, Kashikar N, Hagen V, Krause E, Kaupp UB. A K^+^-selective cGMP-gated ion channel controls chemosensation of sperm. Nat Cell Biol. 2006 Oct;8(10):1149–54. doi: 10.1038/ncb1473. Epub 2006 Sep 10. PMID: 16964244.

Strünker T, Goodwin N, Brenker C, Kashikar ND, Weyand I, Seifert R, Kaupp UB. The CatSper channel mediates progesterone-induced Ca^2+^ influx in human sperm. Nature. 2011 Mar 17;471(7338):382–6. doi: 10.1038/nature09769. PMID: 21412338.

Taiwo BG, Frettsome-Hook RL, Taylor AE, Correia JN, Lefievre L, Publicover SJ, Conner SJ, Kirkman-Brown JC. Complex combined steroid mix of the female tract modulates human sperm. Reprod Biol. 2021 Dec;21(4):100561. doi: 10.1016/j.repbio.2021.100561. Epub 2021 Oct 4. PMID: 34619633.

Tamburrino L, Marchiani S, Muratori M, Luconi M, Baldi E. Progesterone, spermatozoa and reproduction: An updated review. Mol Cell Endocrinol. 2020 Oct 1;516:110952. doi: 10.1016/j.mce.2020.110952. Epub 2020 Jul 24. PMID: 32712385.

Tateno H, Krapf D, Hino T, Sánchez-Cárdenas C, Darszon A, Yanagimachi R, Visconti PE. Ca^2+^ ionophore A23187 can make mouse spermatozoa capable of fertilizing in vitro without activation of cAMP-dependent phosphorylation pathways. Proc Natl Acad Sci U S A. 2013 Nov 12;110(46):18543–8. doi: 10.1073/pnas.1317113110. Epub 2013 Oct 15. PMID: 24128762; PMCID: PMC3831971.

Torres-Flores V, Picazo-Juárez G, Hernández-Rueda Y, Darszon A, González-Martínez MT. Sodium influx induced by external calcium chelation decreases human sperm motility. Hum Reprod. 2011 Oct;26(10):2626–35. doi: 10.1093/humrep/der237. Epub 2011 Aug 2. PMID: 21810864; PMCID: PMC3174032.

Trötschel C, Hamzeh H, Alvarez L, Pascal R, Lavryk F, Bönigk W, Körschen HG, Müller A, Poetsch A, Rennhack A, Gui L, Nicastro D, Strünker T, Seifert R, Kaupp UB. Absolute proteomic quantification reveals design principles of sperm flagellar chemosensation. EMBO J. 2020 Feb 17;39(4):e102723. doi: 10.15252/embj.2019102723. Epub 2019 Dec 27. PMID: 31880004; PMCID: PMC7024835.

Tsien RY, Hladky SB. A quantitative resolution of the spectra of a membrane potential indicator, diS-C3-(5), bound to cell components and to red blood cells. J Membr Biol. 1978 Jan 12;38(1-2):73-97. doi: 10.1007/BF01875163. PMID: 625049.

Vavrusova M, Munk MB, Skibsted LH. Aqueous solubility of calcium L-lactate, calcium D-gluconate, and calcium D-lactobionate: importance of complex formation for solubility increase by hydroxycarboxylate mixtures. J Agric Food Chem. 2013 Aug 28;61(34):8207–14. doi: 10.1021/jf402124n. Epub 2013 Aug 16. PMID: 23906043.

Wang H, McGoldrick LL, Chung JJ. Sperm ion channels and transporters in male fertility and infertility. Nat Rev Urol. 2021 Jan;18(1):46–66. doi: 10.1038/s41585-020-00390-9. Epub 2020 Nov 19. PMID: 33214707; PMCID: PMC7852504.

Williams HL, Mansell S, Alasmari W, Brown SG, Wilson SM, Sutton KA, Miller MR, Lishko PV, Barratt CL, Publicover SJ, Martins da Silva S. Specific loss of CatSper function is sufficient to compromise fertilizing capacity of human spermatozoa. Hum Reprod. 2015 Dec;30(12):2737–46. doi: 10.1093/humrep/dev243. Epub 2015 Oct 8. PMID: 26453676; PMCID: PMC4643530.

Windler F, Bönigk W, Körschen HG, Grahn E, Strünker T, Seifert R, Kaupp UB. The solute carrier SLC9C1 is a Na^+^/H^+^-exchanger gated by an S4-type voltage-sensor and cyclic-nucleotide binding. Nat Commun. 2018 Jul 18;9(1):2809. doi: 10.1038/s41467-018-05253-x. Erratum in: Nat Commun. 2020 Aug 19;11(1):4210. doi: 10.1038/s41467-020-18023-5. PMID: 30022052; PMCID: PMC6052114.

Woodford CR, Frady EP, Smith RS, Morey B, Canzi G, Palida SF, Araneda RC, Kristan WB Jr, Kubiak CP, Miller EW, Tsien RY. Improved PeT molecules for optically sensing voltage in neurons. J Am Chem Soc. 2015 Feb 11;137(5):1817–24. doi: 10.1021/ja510602z. Epub 2015 Jan 29. PMID: 25584688; PMCID: PMC4513930.

Young S, Schiffer C, Wagner A, Patz J, Potapenko A, Herrmann L, Nordhoff V, Pock T, Krallmann C, Stallmeyer B, Röpke A, Kierzek M, Biagioni C, Wang T, Haalck L, Deuster D, Hansen JN, Wachten D, Risse B, Behre HM, Schlatt S, Kliesch S, Tüttelmann F, Brenker C, Strünker T. Human fertilization in vivo and in vitro requires the CatSper channel to initiate sperm hyperactivation. J Clin Invest. 2024 Jan 2;134(1):e173564. doi: 10.1172/JCI173564. PMID: 38165034; PMCID: PMC10760960.

Zeng Y, Clark EN, Florman HM. Sperm membrane potential: hyperpolarization during capacitation regulates zona pellucida-dependent acrosomal secretion. Dev Biol. 1995 Oct;171(2):554–63. doi: 10.1006/dbio.1995.1304. PMID: 7556936.

Zeng XH, Yang C, Kim ST, Lingle CJ, Xia XM. Deletion of the Slo3 gene abolishes alkalization-activated K^+^ current in mouse spermatozoa. Proc Natl Acad Sci U S A. 2011 Apr 5;108(14):5879–84. doi: 10.1073/pnas.1100240108. Epub 2011 Mar 22. PMID: 21427226; PMCID: PMC3078394.

